# Selective uptake into inflamed human intestinal tissue and immune cell targeting by wormlike polymer micelles

**DOI:** 10.1101/2021.01.26.428316

**Authors:** Elena Gardey, Fabian H. Sobotta, Drilon Haziri, Philip C. Grunert, Maren T. Kuchenbrod, Franka V. Gruschwitz, Stephanie Hoeppener, Michael Schumann, Nikolaus Gaßler, Andreas Stallmach, Johannes C. Brendel

## Abstract

Over the 21st century, inflammatory bowel disease (IBD) has become a global disease with no causal therapeutic options. Selective targeting of inflamed areas in the gastrointestinal tract could be an effective treatment circumventing severe side effects for healthy tissue. Our study demonstrates that the shape of polymeric nanostructures represents so far rarely addressed key to required tissue selectivity in the intestine. *Ex vivo* experiments on human colonic biopsies revealed that crosslinked wormlike micelles featuring a dense poly(ethylene oxide) (PEO) shell exclusively enter the inflamed human mucosa without affecting healthy tissue. Similarly designed spherical micelles (∼25 nm) or vesicles (∼120 nm) penetrate both tissues or were barely uptaken at all, respectively. Moreover, it was found that the particles colocalize with immune cells in the lamina propria facilitating a specific targeting of the main pro-inflammatory cells within the diseased human mucosa. These findings demonstrate an untapped potential in particle design and enable new vistas for an effective treatment of IBD.

---

More than six million people worldwide suffer from inflammatory bowel disease (IBD) and numbers are rising which might pose a substantial burden to global health care systems^1-3^. Current treatments rely on anti-inflammatory drugs such as steroids or immunosuppressants including cytokine antibodies which either cannot adequately reduce the symptoms or induce severe side effects^4-6^. In the last decades research on nano/microparticles have attracted considerable attention promising local targeting of inflamed sites^7-9^ by inducing an increased permeability and disruption of the local intestinal barrier at the luminal side^6,10-12^. As a consequence, tremendous effort has been paid to optimize these nanoparticle systems for improve their selective uptake into inflamed areas. Targeting strategies comprise the adjustment of surface charges, the selective local degradation, or receptor mediated targeting^6,13,14^. For example, the surface modification with poly(ethylene oxide) (PEO) appears to enhance the penetration into inflamed tissue through the epithelium^15,16^, as an interaction with the mucus is suppressed^15,17^ if the surface of the particles is densely covered^16^. A rather straightforward method to increase tissue selectivity is achievable by adjusting the size of the micro/nanoparticles and several studies confirmed an accumulation of nano/micrometer sized materials at inflamed sites^10,11^.

Despite these promising developments, a major challenge remains to maintain the desired high selectivity for inflamed areas and targeting of macrophages as main pro-inflammatory cells in these areas^18^. Interestingly, the factor shape has so far barely been investigated for polymer-based nanostructures, despite the previously mentioned importance of particle dimensions. Its explicit impact on the particle interaction with intestinal barriers becomes apparent in pioneering studies on co-culture models of intestinal epithelium layers^19^. Only very recently, the therapeutic advantage of one-dimensional nanotubes based on assembled peptides was demonstrated *in vivo* on dextran sulfate sodium induced ulcerative colitis^20^. The limited number of studies is surprising considering the beneficial effects of soft elongated nanostructures reported for crossing other biological barriers or specific targeting^19-25^. In an interdisciplinary attempt we combined our expertise on creating polymer nanostructures of various shapes and *ex vivo* testing of particulate systems on human colonic macrobiopsies to evaluate specifically the impact of shape in case of soft polymer nanostructures. We hypothesized that the anisotropic shape and soft nature of wormlike polymer particles enables an efficient translocation into inflamed intestinal tissue based on the nanometer sized diameter, while the length in the micrometer scale prevents undesired uptake through healthy epithelial cell layers. To circumvent previously observed discrepancies between animal studies and clinical tests on humans^8^, we focused in our study on *ex vivo* experiments using biopsies obtained from healthy patients without inflammation and patients with IBD^11,15^. Mounted into Ussing chambers the translocation and accumulation of nanoparticles within the human tissue can be monitored and the local distribution can be analyzed by fluorescence microscopy of tissue cross-sections.

The required nanoparticles of different shape were prepared by self-assembly of amphiphilic block copolymers into micelles and subsequent crosslinking. Such polymeric micelles have already demonstrated their potential in nanomedicine^26-29^. Advantages of these self-assembled materials lie in the access to very small structures (< 50 nm) while maintaining narrow size distributions and the uniform and dense packing of hydrophilic chains on the surface (e.g. PEO). Another benefit of the assembly process is the ability to form nanostructures of different shapes including spherical micelles^30^, wormlike micelles (filomicelles)^31^ and polymeric vesicles (polymersomes)^32^. These shape variations are usually achieved by adjusting the polymer composition, but the preparation conditions can also play a crucial role in the self-assembly^33-36^. Here we take advantage of the latter and by scrutinizing different methods we identified suitable conditions to create all three above mentioned shapes from the same polymer. As a consequence, any influence of polymer nature and composition is eradicated and solely the shape of the nanostructures impacts the uptake behavior.

## Polymer synthesis and formation of different nanostructures

The polymer micelles were formed from an amphiphilic block copolymer which consists of a hydrophobic butyl acrylate copolymer (PBA) and PEO as hydrophilic block. 10 mol% of pyridyldisulfide ethyl acrylate (PDSA) were further included in the hydrophobic block to enable a sufficient crosslinking of the micelles after their assembly, which was found to minimize potential pro-inflammatory effects of such nanostructures^27,37,38^. The copolymer of PBA and PDSA was synthesized by reversible addition−fragmentation chain-transfer (RAFT) polymerization to guarantee a narrow molar mass distribution of the polymer chains (see SI for experimental details and characterization; Supplementary Table 1, Fig. 1, 2). The obtained polymers were then coupled to a heterotelechelic α-amino/ω-azido PEO (5 kDa) *via* amide formation using an activated ester approach to finally yield a well-defined (dispersity Ð < 1.15) amphiphilic block copolymer P(BA-*co*-PDSA)-*b*-PEO.

**Fig 1.**
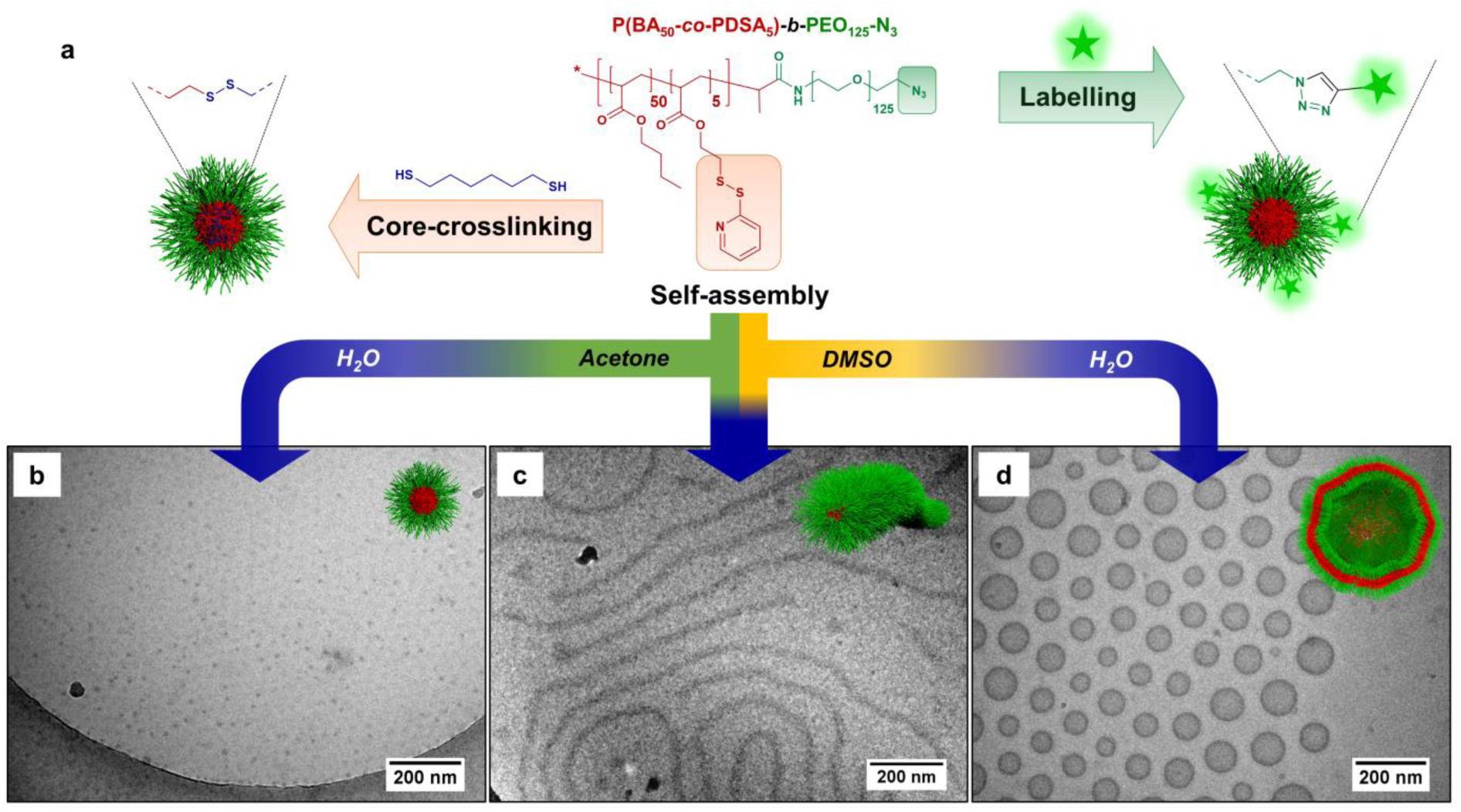
Preparation of core-crosslinked polymeric nanostructures with different shapes based on the same polymer. **a**, Schematic representation of the solvent switch approach to generate nanostructures of different morphology including subsequent core-crosslinking and labeling steps. **b**, cryo-TEM image of spheres **M1-CL\*** prepared with acetone as co-solvent after crosslinking and labeling. **c**, cryo-TEM of worm-like micelles **M3-CL\*** prepared with DMSO/acetone as co-solvent after crosslinking and labeling. **d**, cryo-TEM of vesicles **M2-CL\*** with DMSO as co-solvent after crosslinking and labeling.

**Fig 2.**
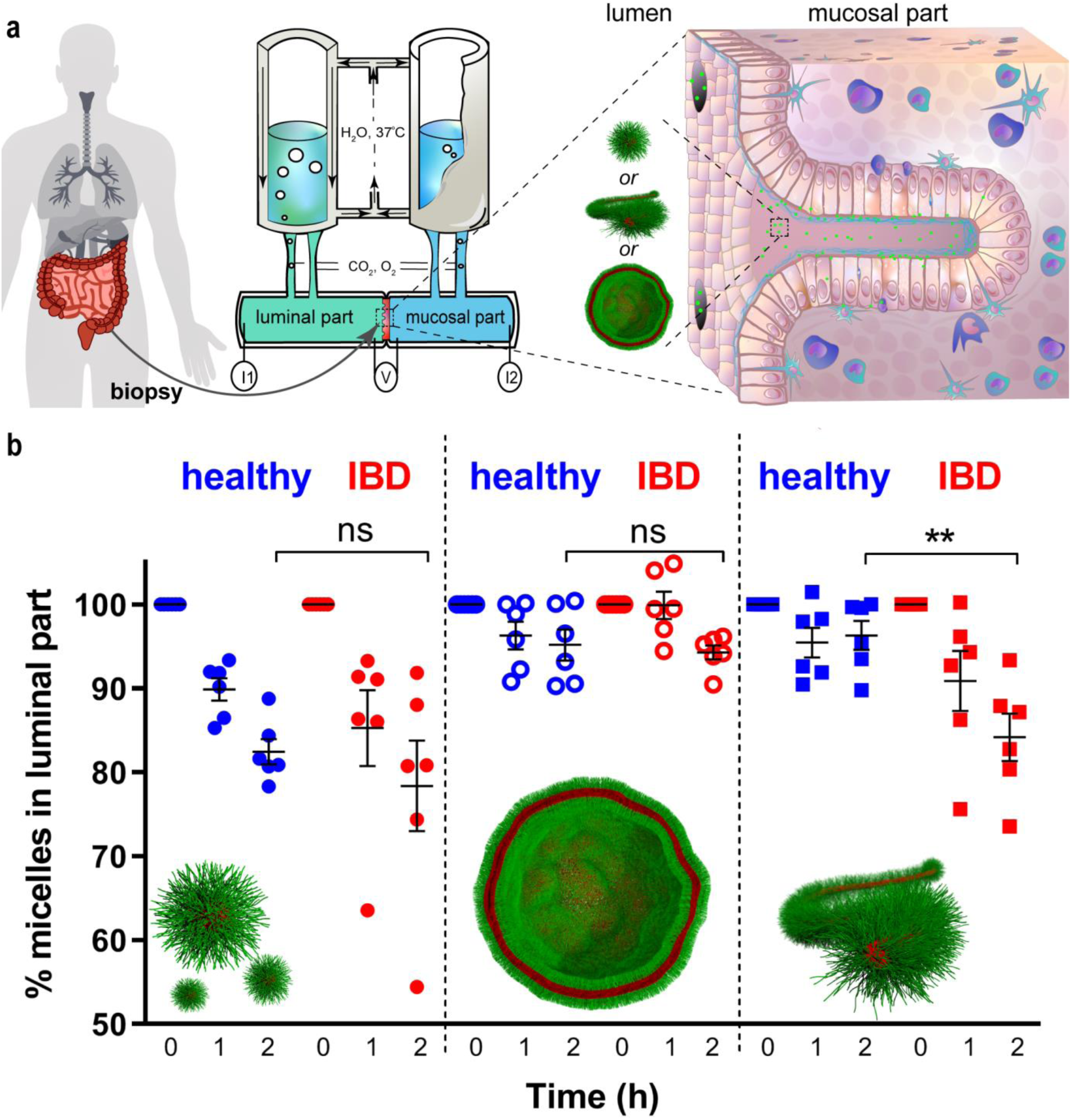
*Ex vivo* experiment on nanoparticle interaction with healthy and inflamed human mucosa. **a**, Schematic representation of the experimental setup using Ussing chambers with mounted biopsies. Nanoparticles (represented by green dots) were added to the luminal part of the Ussing chambers. The modified Krebs-Ringer bicarbonate buffer in the Ussing chamber is continuously oxygenated and kept at 37°C. Integrity and viability of the biopsy are controlled via inserted electrodes (V, I). **b**, Time-dependent change of the concentration of spherical micelles (full dots), polymeric vesicles (empty dots), and worms (squares) in the luminal part with healthy (blue) and inflamed (red) tissue. The exposed tissue area is 4.9 mm^2^. The results are presented as percentage of the initial concentration (M ± SEM); ns -not significant, **p < 0.001, n = 6 (n – number of different donors/patients in each group). Two-tailed unpaired Student’s t-test was used. IBD -inflammatory bowel disease.

The self-assembly of these block copolymers into different micelles was subsequently induced by a solvent switch from a good solvent for both blocks (DMSO or acetone) to the selective solvent water (the general approach is depicted in Fig. 1).

Therefore, water was slowly added to a solution of the polymer until a volume ratio of 1:1 of the solvents was reached, which already triggered the particle formation. To stabilize the obtained micellar morphologies, 1,6-hexane dithiol was added as a crosslinker to form stable nanoparticles. This compound diffuses into the micellar core and reacts with the PDSA units to form disulfide links. The reaction can conveniently be monitored by the increasing absorbance of the formed by-product 2-mercaptopyridine (Supplementary Fig. 3). The successful crosslinking was additionally verified by dissolution experiments in the non-selective solvent THF where the nanoparticle structure is maintained after the reaction (Supplementary Fig. 4). Furthermore, no changes in the hydrodynamic diameter are observed in DLS (Supplementary Table 2, Fig. 5). The residual solvent was finally removed by dialysis.

Interestingly, a substantial difference in the resulting morphologies of the assemblies was observed depending on the initial solvent. Rather small nanoparticles with a hydrodynamic diameter *D*_H_ of ∼ 25 nm and low dispersity (Supplementary Table 3, Fig. 6a,b), were formed when starting from polymers dissolved in acetone. Characterization by cryogenic transmission electron microscopy (cryo-TEM) confirmed the presence of spherical shaped micelles (Fig.1b). On the other hand, replacing acetone with DMSO as initial solvents induces a complete phase transition towards vesicles or polymersomes (Figure 1d). Their diameter is significantly larger (*D*_H_ ∼ 116 nm) but a narrow size distribution (PDI < 0.05) was maintained (Supplementary Table 3, Fig. 6a,b). Hence, the choice of the solvent seems to substantially impact the assembly and enables reproducible access to commonly opposing solution morphologies in block copolymer self-assembly^39-41^. Assuming divergent surface effects and assembly kinetics in the different solvents, we tested also mixtures of the two solvents acetone and DMSO. Analogous to typical phase transitions of block copolymer solutions, the 1:1 mixture of the solvents led to long wormlike micelles or filomicelles (Fig. 1c, Supplementary Table 2, 3, Fig. 6a,b). The cross-sectional diameter of the worms (*D*_TEM_ = 20.5 nm) was comparable to the *D*_TEM_ of the spherical micelles (*D*_TEM_ = 15.5 nm), while their length varied between150 nm to 2 µm. Additional analysis by asymmetric flow-field flow fractionation (AF4) displayed clear shifts in the elution time with surprisingly narrow distributions, which verifies the presence of distinct morphologies in each sample (Supplementary Fig. 6, 7). It is noteworthy that such full transitions between different morphologies usually require a variation of the composition of the polymer building blocks and similar sharp transitions by process modifications remain exceptional. In our case, all three main morphologies (spheres, worms, and vesicles) can selectively be targeted from the same block copolymer.

For localization of the nanoparticles in the biological experiments, the nanoparticles were finally labelled with the fluorescent dye AF488 using strain-promoted azide-alkyne coupling (see SI for further details). The successful conjugation and the absence of unbound dye after purification were confirmed by size exclusion chromatography with UV-vis detection (Supplementary Fig. 2b).

## Ex vivo uptake and translocation of nanoparticles by healthy and inflamed human mucosa

With these particles of different shape at hand, we focussed on the evaluation of translocation efficiency through the epithelial barrier and uptake of these polymeric nanoparticles into human mucosa. Therefore, biopsies of inflamed tissue from patients with IBD were mounted in Ussing chambers (Fig. 2a) that allow controlling the tissue viability and integrity in *ex vivo* experiments (see Supplementary for experimental details on the procedure). Healthy individuals undergoing screening colonoscopy served as controls.

After confirming the correct placement and integrity of the tissue, equal amounts of polymeric nanoparticles (final concentration: 100 µg mL^-1^) were injected at the luminal side of the tissue in the Ussing chambers. Any change in concentration of the applied nanoparticles was monitored over 2 h by taking aliquots from the solutions on the two sides of the biopsies – luminal and mucosal part – and measuring the respective fluorescent intensity (Fig. 2).

Small spherical micelles (∼ 25 nm) appear to significantly enter both tissues (*p* < 0.0001, *p* = 0.0057, Supplementary Table 5) as a continuous decrease of the applied concentration in the luminal side is observed over 2 h. Although a slightly faster rate seems to apply in case of inflammation (concentration reduction of 21.6% ± 5.4 after 2h), no significant difference (*p* = 0.4847) was observed to healthy tissue (17.6% ± 1.5 after 2h), which is similarly penetrated by such small particles. Interestingly, only an insignificant translocation (< 0.5% for inflamed tissue) to the mucosal side is observed in these cases (Supplementary Fig. 8a,b), which implies that the reduced amount in the luminal part must be related to a particle accumulation in the tissue (control experiments without tissue rebutted any significant secondary losses by interaction with the chambers, Supplementary Fig. 9). Furthermore, only insignificant amounts (1 to 1.5%, see Supplementary Fig. 8d) of the particles could be washed off the surface of the biopsies after unmounting the samples, which support an uptake into the tissue.

The limited difference between inflamed and healthy tissue is surprising, considering a previously often reported selectivity of nanoparticles, but it must be considered that most examined particles still feature sizes > 100 nm. The tested polymeric vesicles (∼120 nm) are therefore of a more comparable size to evaluate this effect. However, in this case the uptake is considerably reduced and only a minor reduction of the concentration (5.8% ± 0.9, *p* = 0.002, Supplementary Table 5) was observed for inflamed biopsies after 2 h (Fig. 2). Although there is an increase in uptake compared to 1 h, which, the difference to healthy mucosa was found to be not statistically significant. Similar to the small spherical micelles only minor amounts translocated to the mucosal side in case of the inflamed tissue. This discrepancy to literature reports can only be related to the difference in the nature of the particles, as the tested vesicles should be softer than common dense polymer nanoparticles based on poly(lactide-*co*-glycolic acid) (PLGA) and feature a very dense layer of PEO on their surface. The most distinct differences between healthy and inflamed tissue were observed, when crosslinked filomicelles were applied. The experiment on inflamed tissue from patients with IBD confirmed a comparably fast uptake (15.9% ± 2.8 after 2 h) as for the small spherical micelles with similarly low translocation to the mucosal part. However, only insignificant amounts of filomicelles entered the healthy biopsies. The filomicelles therefore offer an excellent selectivity for targeting inflamed tissue with clear statistical difference after 2 h (*p* = 0.0041).

It has to be mentioned that the uptake rate by human colon mucosa not only depends on size and structure of the applied nanoparticles but depends moreover on the degree of inflammation in the tissue and the severeness of structural changes in the intestinal barriers. In our study we used only biopsies from IBD patients without severe inflammation and tissue damage (Supplementary Table 4). In this context, the observed selectivity of filomicelles is even more remarkable.

## Localisation of the nanoparticles in human mucosa

After the Ussing chamber experiments, the biopsies were further analyzed to investigate the localisation of the nanoparticles within the tissue. Therefore, the samples were cryosectioned into thin slices (see Fig. 3a for an illustration) and fluorescence microscopy was used to locate the labelled particles within the lamina propria of healthy and inflamed tissue (Fig. 3b).

**Fig 3.**
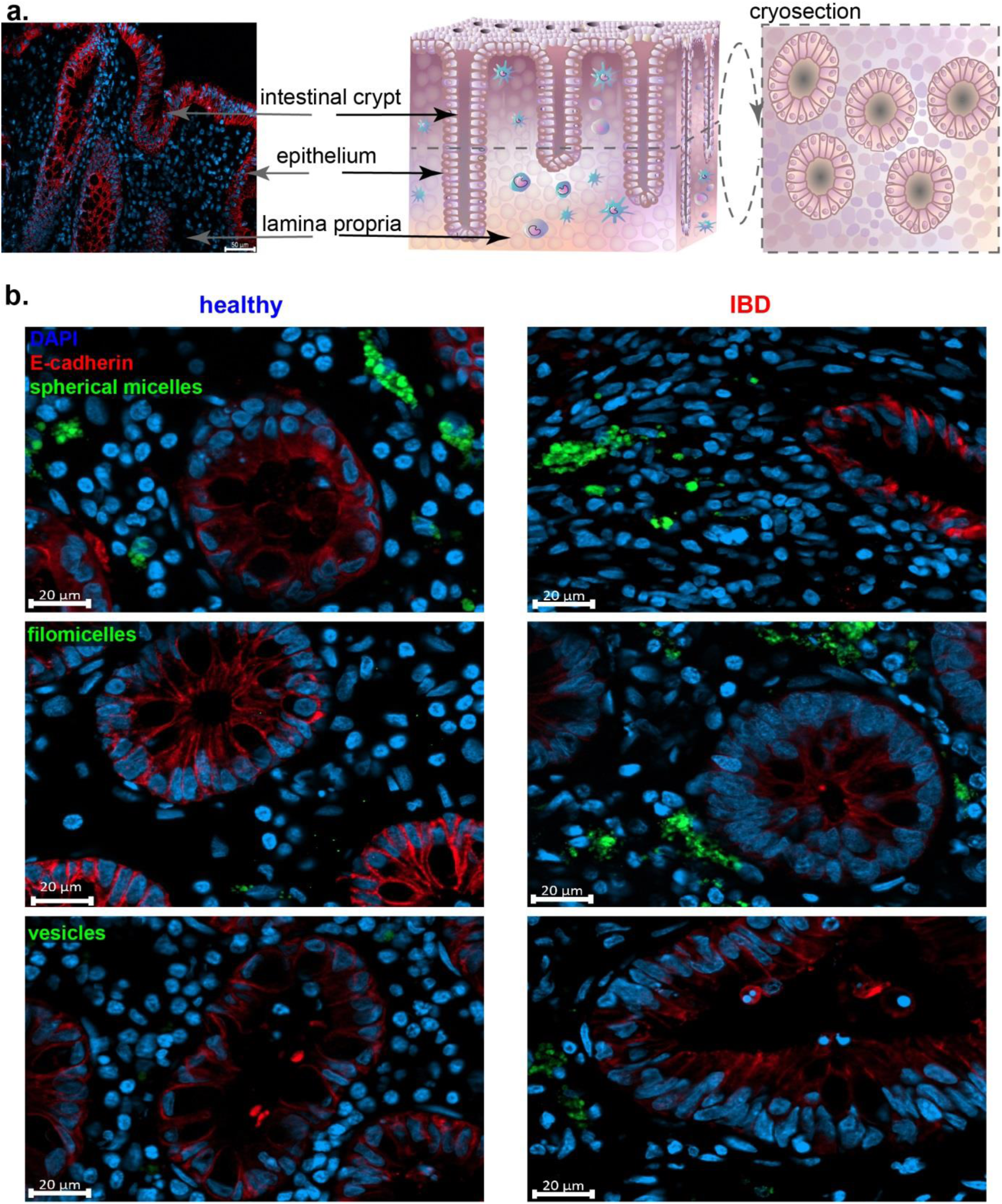
Localisation of nanoparticles in healthy and inflamed human mucosa. **a**, The scheme of colon mucosa demonstrates lamina propria with immune cells, epithelial cells, intestinal crypts and the orientation of cryosection. **b**, Localisation of spherical micelles, wormlike micelles and vesicles in human mucosa after 2 h incubation in Ussing chambers (green: nanoparticles; blue (DAPI): nuclei; red: E-cadherin (epithelium)). Biopsies were sectioned (6 µm) by cryotome slicing. Scale bar is 20 µm.

A comparison of the microscopy images confirms our initial study on the uptake. The small spherical particles appear distributed throughout both healthy and inflamed tissues, with comparable intensities. Wormlike micelles, however, are only observed in the lamina propria of inflamed tissue in significant amounts, while barely any particles can be found in the biopsies of healthy controls. The larger vesicles did result in only few signals in both tissues, which is in agreement with their insignificant uptake. These trends are confirmed in a more comprehensive analysis of z-stacks over a thickness of 6 µm (see Supplementary Videos 1-6 for more details).

Interestingly, not all cells of the lamina propria appear to uptake the nanoparticles in similar amounts. Clear accumulation of the labelled micelles in few cells can be observed in the microscopy images (Fig. 4). Since we hypothesize that the nanoparticles are mainly uptaken by immune cells of the lamina propria, we co-stained the slices of the biopsies with the CD11b marker that selectively adheres to immune cells, including macrophages, dendritic cells, monocytes, neutrophils and natural killer cells^42-44^.

**Fig 4.**
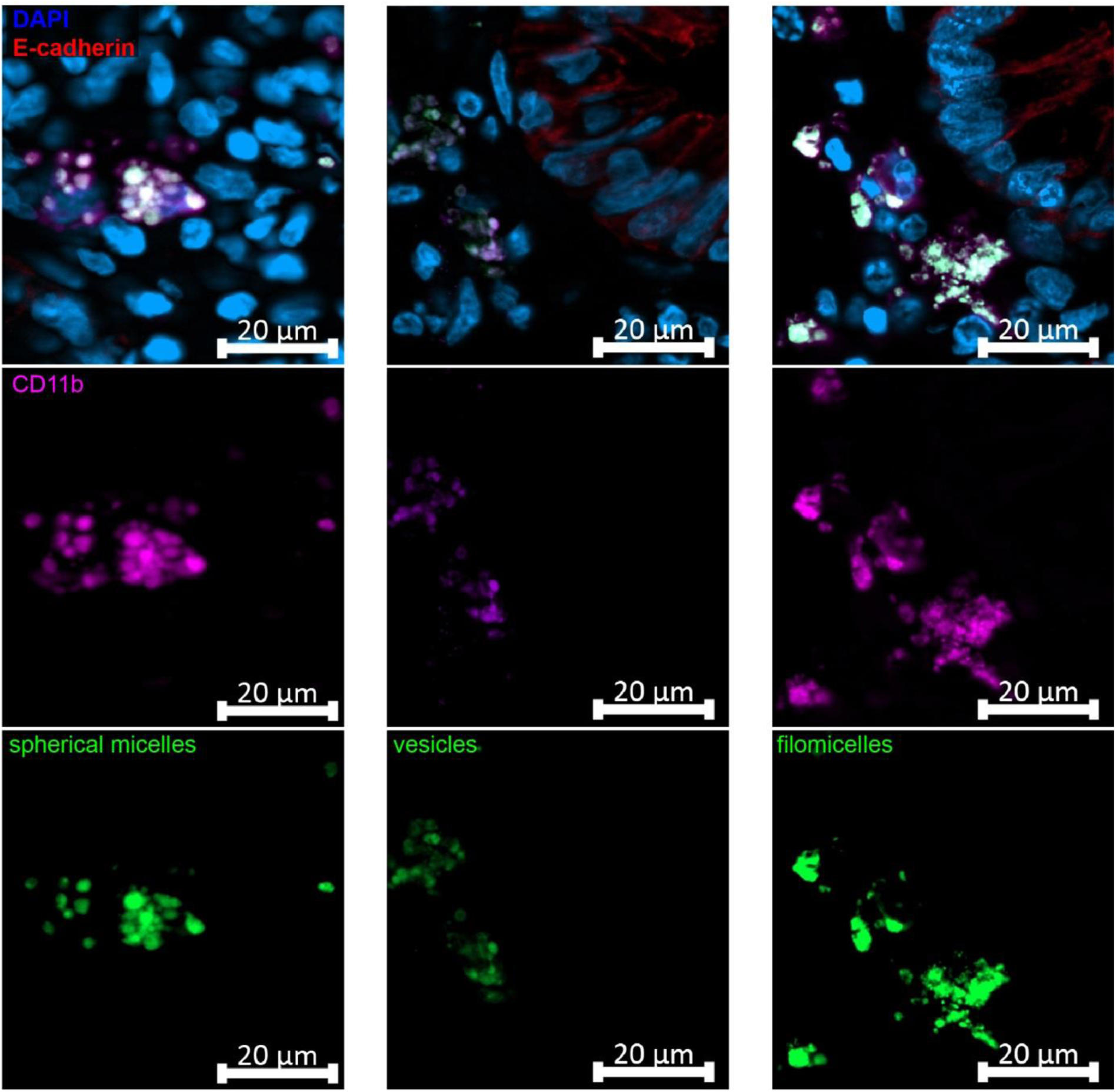
Accumulation of micelles in immune cells of inflamed human mucosa. Accumulation of spherical micelles, vesicles and wormlike micelles in immune cells CD11b (violet), green: nanoparticles; blue (DAPI): nuclei; red: E-cadherin (epithelium). Biopsies were sectioned (6 µm) by cryotome slicing. Scale bar is 20 µm.

This immunohistochemistry revealed a clear overlap with the applied polymer micelles and thus confirms our hypothesis of a selective accumulation in immune cells of the lamina propria (Fig. 4; Supplementary Fig. 10-12). In addition, this more detailed analysis of the images revealed a considerably increased number of immune cells within the tissue from IBD patients. Nevertheless, the distinct selectivity for uptake into these immune cells further underlines the potential of these polymeric micelles to target the most important pro-inflammatory cells within the lamina propria. Similar selectivity usually requires active targeting of cell specific receptors^45,46^. In accordance with previous studies, we also observed a disintegration of epithelium in biopsies from IBD patients which is indicated by their variable E-cadherin staining ability. Disturbing the integrity of the epithelium has already been accounted for size-dependent uptake of nanoparticles and we also rationalize our observed tissue selectivity by the enhanced permeability^28,47^.

## Conclusion

The ability to create different micelle morphologies from the same amphiphilic block copolymer represents a unique possibility for the exclusive analysis of the impact of shape on the uptake into human mucosa of healthy and inflamed tissue. We here demonstrate that a PEO-based blockcopolymer comprising a hydrophobic block (PBA), which can be crosslinked to form stable nanostructures, can selectively and exclusively be assembled into spherical micelles, vesicles and wormlike micelles (filomicelles). Our subsequent study on such densely PEO-covered nanoparticles reveals that their penetration through the gastrointestinal barrier unequivocally depends on their size and the geometry. Localization of nanoparticles in healthy and inflamed mucosa proves a beneficial behaviour of filomicelles compared with spherical micelles and vesicles. Flexibility notwithstanding, the wormlike micelles cannot penetrate through the healthy gastrointestinal barrier, while they easily translocate across the disturbed epithelium of inflamed tissue in similar rates as much smaller spherical micelles. The rather low diameter must play a key role in this context, as vesicles of a larger diameter are not able to cross the disturbed barrier of inflamed tissue in significant amounts. In combination with the observed passive targeting of immune cells (the principal aim for treatment in IBD^18^) in the mucosa of patients with IBD, the presented wormlike micelles represent an innovative tool for efficient and exclusive targeting of inflamed areas and the residing main proinflammatory cells in case of IBD. The comparably high volume of these wormlike nanostructures compared to spherical nanoparticles further promises an effective transport of anti-inflammatory drugs to these targets^48^, which is currently under investigation. The possibility to further modify the large surface area of these nanostructures bears moreover enormous potential to expand the selectivity and further increase their accumulation within immune cells of the inflamed mucosa. Additional impact will certainly be provided by the adjustment of the length of the filomicelles, which is part of an ongoing study.

Overall, we conclude that the shape of polymeric nanostructures represents a key factor for selective treatment of inflamed areas in the colon, which certainly deserves more attention in the future design of effective treatments of IBD. In this regard, emphasis should also be paid in future to the development of straightforward preparation techniques to increase the accessibility of wormlike nanostructures to reveal their full potential.

## Acknowledgements

This work was financially supported by the DFG-funded Collaborative Research Centre PolyTarget (Project-ID: 316213987 – SFB 1278; Project A05, Z01). Johannes C. Brendel further thanks the DFG for funding within the Emmy-Noether Programme (Project-ID: 358263073) and support by the FCI (Fonds der Chemischen Industrie). We thank D. Cardoso da Silva (Charité-University Medicine, Berlin) and Dr. F. Dengler (University of Leipzig) for consultation on Ussing chamber equipment. We thank all doctors of the endoscopy department for their contribution, especially M. Kolleck and Dr. med. U. Mühlenberg. Prof. U. S. Schubert is furthermore acknowledged for his continuous support and access to excellent research facilities. Cryo-TEM investigations were conducted on the Jena Center for Soft Matter (JCSM) electron microscopy facilities established with funds by the DFG and the European Funds for Regional development (EFRE).

## Author contributions

E.G. performed the Ussing chamber experiments, cryosectioning, IHC, fluorescent microscopy, wrote the paper. F.H.S. performed the synthesis and characterization of nanoparticles, wrote the paper. D.H. and P.C.G organized the human material, selected suitable healthy donors and patients with IBD. F.V.G. performed the AF4 measurements. M.T.K. and S.H. performed the cryo-TEM imaging. M.S. consulted in establishing of the Ussing chamber experiments and handling of the human biopsies. N.G. evaluated the severeness of the inflammation in IBD patients. A.S. and J.C.B. supervised the project, wrote the paper. All authors edited and approved the final manuscript.

## Supplementary information

Methods and additional data are presented in Supplementary information.

