## Supplementary Information for "Selective uptake into inflamed human intestinal tissue and immune cell targeting by wormlike polymer micelles"

### *Materials*

All chemicals and solvents were purchased from Sigma-Aldrich, Acros Organics, Carl Roth, Merck, Jena Bioscience and were used without further purification unless mentioned otherwise. Pyridyldisulfide ethyl acrylate (PDSA) and 2-(Butylthiocarbonothioylthio)propanoic hydroxysuccinimide (PABTC-NHS) were prepared *via* previously reported procedures[1,2]. 1,4-dioxane and butyl acrylate (BA) were treated 24 h with inhibitor remover resin prior to use. $N_3$-$PEO_{125}$-$NH_2$ was purchased from Rapp Polymere.

### *Characterization methods*

*Nuclear magnetic resonance (NMR) measurements*

$^1$H-NMR was performed at room temperature on a Bruker AC 300 MHz spectrometer.

*Size-exclusion chromatography (SEC)*

The polymers were analyzed on a Shimadzu system equipped with a SCL-10A system controller, a LC-10AD pump, a RID-10A refractive index detector and a PSS SDV column with *N*,*N*-dimethylacetamide (DMAc) + 0.21% LiCl as eluent. The column oven was set to 50 °C.

*Asymmetrical flow field-flow fractionation (AF4)*

AF4 measurements were performed on an AF2000 MT System from Postnova Analytics GmbH Landsberg, Germany), equipped with a tip and focus pump (PN1130), an autosampler (PN5300), and a channel oven unit (PN4020) set to 25°C. The channel was coupled to a multiangle laser light scattering (MALLS) detector (PN3621) equipped with a 532 nm laser and measuring 21 angles, a refractive index (RI) detector (PN3150), and a UV-detector (PN3212) set to 280 nm. The channel had a trapezoidal geometry with a nominal height of 350 µm. Regenerated cellulose (RC) membrane from Postnova Analytics GmbH (10 kDa RC membrane) with a molar mass cutoff of 10 kDa was used as accumulation wall. As the mobile phase aqueous solution with 0.002 w% of $NaN_3$ was used. 50 µL of the sample at a concentration of 1 mg $mL^{-1}$ was injected with an injection flow rate of 0.2 mL $min^{-1}$, a focus flow rate of 0.8 mL $min^{-1}$, and a cross-flow rate of 0.7 mL $min^{-1}$, resulting in a detector flow rate of 0.3 mL $min^{-1}$. The focusing time was 4 min before switching to elution at an exponentially decaying crossflow from 0.7 mL $min^{-1}$ to 0.2 mL $min^{-1}$ in 76.2 min. Thereafter the crossflow profile was set to decay in a linear way from 0.05 mL $min^{-1}$ to 0.04 mL $min^{-1}$ in 71 min (Supplementary Fig. 1). Before the start of the next measurement, a rinsing step was performed at 1.5 mL $min^{-1}$ flow of the tip pump for 20 min. After each sample measurement, a blank measurement was run which was subtracted from the data of the sample measurement for analysis. The MALLS data of the scattering angles from 20°-148° was analysed via ZIMM plot to obtain the radius of gyration ($R_g$) at the specified elution times.

*Dynamic light scattering (DLS)*

DLS was performed on a ZetaSizer Nano ZS (Malvern, Herrenberg, Germany) equipped with a He–Ne laser operating at a wavelength of λ = 633 nm. Counts were detected at an angle of 173°. The particle size was approximated as the effective diameter (Z-average) obtained by the cumulants method assuming a spherical shape. All measurements were conducted at 25 °C in semi-micro cuvettes after equilibration times of 30 s in triplicate. Every measurement included 10 runs, in which every run took 30 seconds. Apparent hydrodynamic radii were calculated using the Stokes-Einstein Equation (1):

$$R_{\mathrm{h}} = \frac{kT}{6\pi\eta D} \qquad (1)$$

$R_h$ = hydrodynamic radius, k = Boltzmann constant, T = absolute temperature, η = viscosity of the sample and D = apparent translational diffusion coefficient.

*Cryogenic transmission electron microscopy (cryoTEM)*

The measurements were performed on a FEI Tecnai $G^2$ 20 equippedwith a $LaB_6$ filament with anacceleration voltage of 200 kV. Samples were prepared on Quantifoil grids (R2/2) which were treated with Ar plasma prior to use for hydrophilization and cleaning. 8.5 µL of the solution were vitrified on Quantifoil grids using a Vitrobot Mark IV system. Liquid ethane was used as a cryogen. Samples were transferred to a Gatan 626 cryo holder and were maintained at a temperature < -175 °C during the entire process. All images were acquired with a Mega View (OSIS, Olympus Soft Imaging Systems) or an Eagle 4k CCD camera, respectively.

#### *Polymer synthesis*

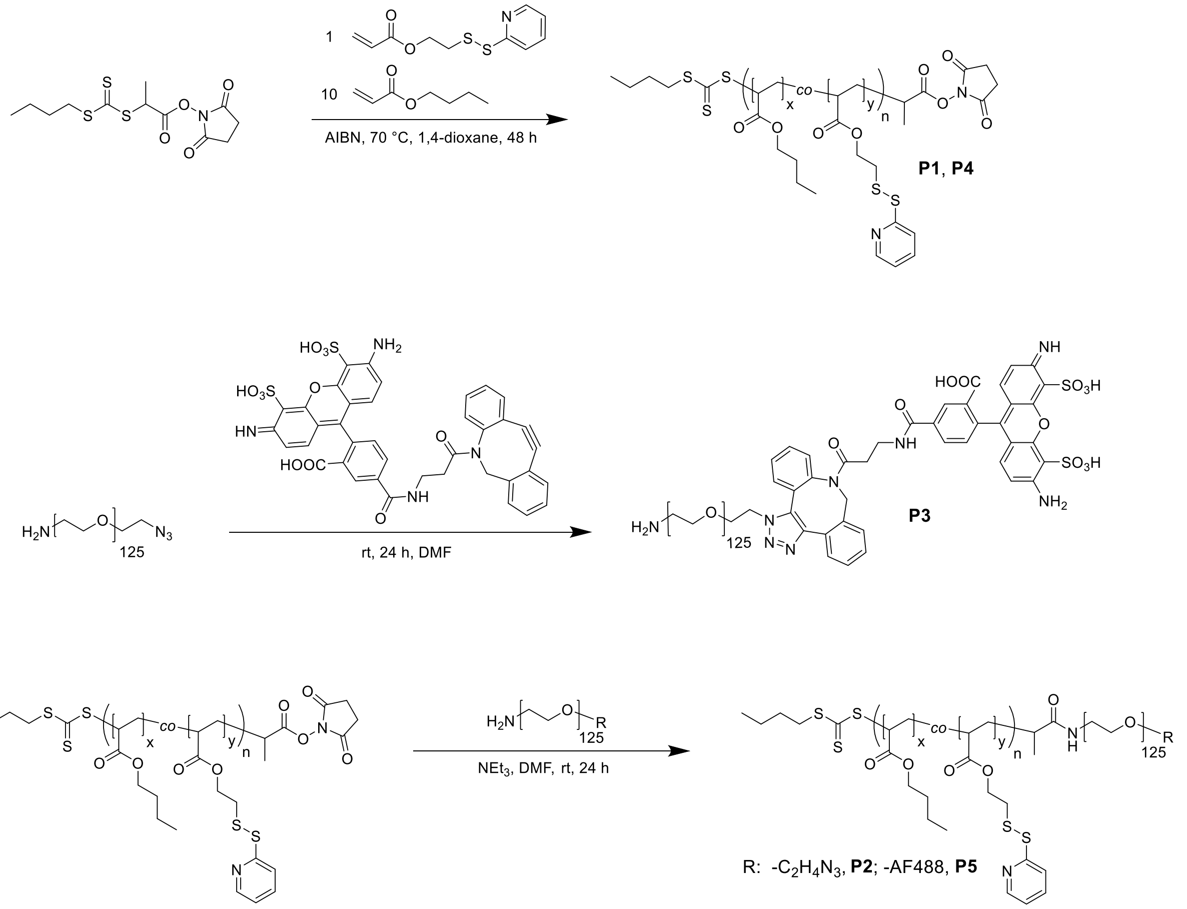


**Scheme 1. Overview of the synthetic steps of the polymer preparation.**

The applied amphiphilic block copolymer is composed of a copolymer of butyl acrylate (BA) and pyridyldisulfide ethyl acrylate (PDSA) as hydrophobic block and PEO as hydrophilic block (Supplementary Scheme 1). In a first step, the copolymer of PBA and PDSA was synthesized by reversible addition−fragmentation chain-transfer (RAFT) polymerization using PABTC-NHS as functionalized chain transfer agent (CTA) for subsequent coupling reaction. BA and PDSA were mixed in a molar ratio of 10 to 1 and polymerized in 1,4-dioxane as solvent. In the next step, the NHS-ester functionalized polymer P($BA_{50}$-*co*-$PDSA_5$)-NHS (**P1**) could be directly coupled to amino end-functionalized PEO ($N_3$-$PEO_{125}$-$NH_2$) of 5 kDa in quantitative yield. The obtained block copolymer P($BA_{50}$-*co*-$PDSA_5$)-*b*-$PEO_{125}$-$N_3$ (**P2**) had an overall molar mass of 15 kDa and a narrow dispersity (Đ < 1.15) after purification by precipitation. All polymers were characterized by $^1$H-NMR and SEC (Supplementary Table 1, Fig. 1, 2a).

**Table 1. Overview of the synthesized polymers including abbreviations and characterization.**

| *ID* | *Polymer* | $M_{n,theo}$[a] [kDa] | $M_{n,SEC}$[b] [kDa] | Đ[b] | *Conversion [%]* | $DP_{BA}$[c] | $DP_{PDSA}$[c] | $DP_{PEO}$ |
|---|---|---|---|---|---|---|---|---|
| ***P1*** | P($BA_{50}$-*co*-$PDSA_5$)-NHS | 8.1 | 9.6 | 1.19 | 79 | 50.3 | 5.3 | - |
| ***P2*** | P($BA_{50}$-*co*-$PDSA_5$)-*b*-$PEO_{125}$-$N_3$ | 13.5 | 18.3 | 1.14 | > 99 | 50.3 | 5.3 | 125 |
| ***P3*** | $H_2N$-$PEO_{125}$-AF488 | 6.3 | 5.4 | 1.07 | > 99 | - | - | 125 |
| ***P4*** | P($BA_{47}$-*co*-$PDSA_5$)-NHS | 7.4 | 8.1 | 1.13 | 73 | 46.5 | 4.7 | - |
| ***P5*** | P($BA_{47}$-*co*-$PDSA_5$)-*b*-$PEO_{125}$-AF488 | 13.7 | 15.0 | 1.11 | > 99 | 46.5 | 4.7 | 125 |

**a,** Calculated based on $[M]_0/[CTA]_0$ × monomer conversion. **b,** Determined by SEC (Eluent: DMAc + 0.21 wt% LiCl, PMMA-calibration). **c,** Calculated as follows: $DP_{BA}$ = (conversion × ([M]/[CTA])) × [BA]/([BA]+[PDSA]), $DP_{PDSA}$ = (conversion × ([M]/[CTA])) × [PDSA]/([BA]+[PDSA]).

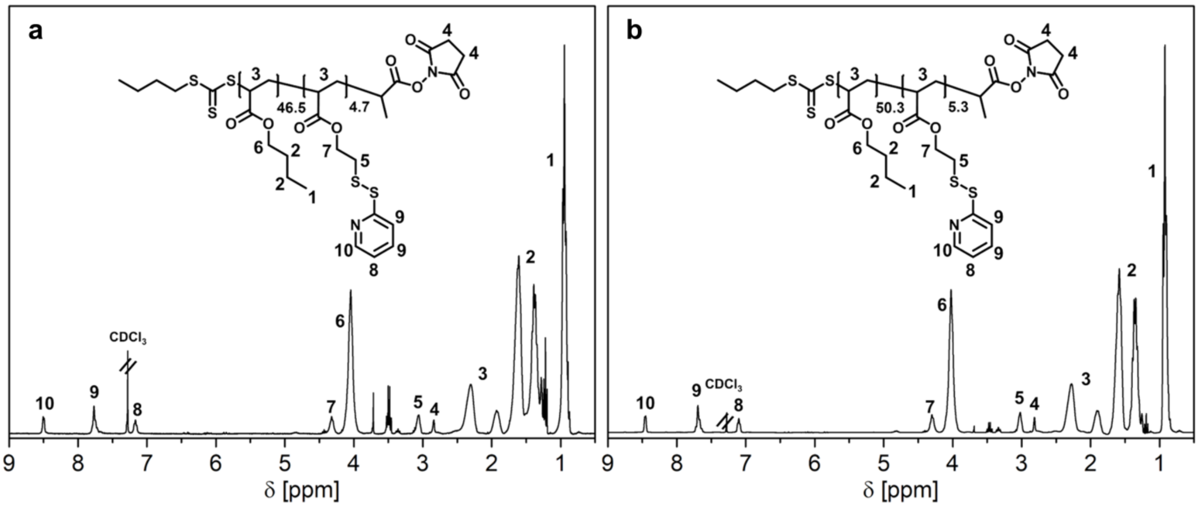

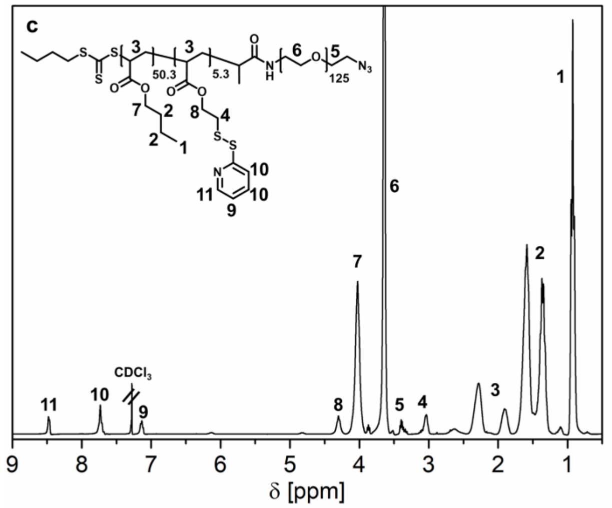


**Fig. 1 Overview of the $^1$H-NMR (300 MHz) spectra measured for polymers after purification. a,** P(BA$_{47}$-*co*-PDSA$_7$)-NHS **P4** in $CDCl_3$. **b,** P(BA$_{50}$-*co*-PDSA$_5$)-NHS **P1** in $CDCl_3$. **c,** (BA$_{50}$-*co*-PDSA$_5$)-*b*-PEO$_{125}$-N$_3$ **P2** in $CDCl_3$.

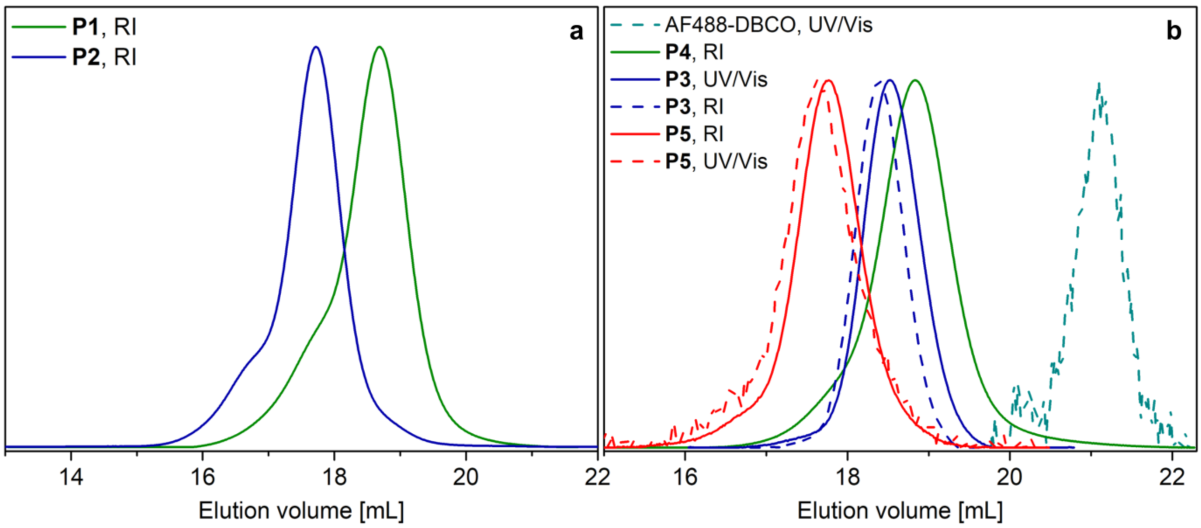


**Fig. 2 Characterization of the synthesized polymers by SEC. a,** Overlay of the SEC curves of the samples **P1** and **P2**. **b,** SEC comparison of fluorescent labeled polymers **P3** and **P5**. (Eluent: DMAc + 0.21% LiCl, PMMA-calibration, UV/Vis detection 480-513 nm).

The dye was conjugated *via* strain-promoted azide-alkyne cycloaddition (SPAAC) between dibenzocyclooctin functionalized AF488 and the azide moiety on the PEO. Two strategies were pursued in this regard: In a first attempt, a fully labelled block copolymer was prepared separately. Therefore, the dye was first attached to N$_3$-PEO$_{125}$-NH$_2$ to give the labelled polymer AF488-PEO$_{125}$-NH$_2$ (**P3**). Subsequently this polymer was coupled to a new batch (comparable composition, see Supplementary Table 1 for details) of the hydrophobic copolymer (P(BA$_{47}$-*co*-PDSA$_5$)-NHS, **P4**) for the formation of a fully labelled block copolymer (P(BA$_{50}$-*co*-PDSA$_5$)-*b*-PEO$_{125}$-AF488, **P5**). This procedure allowed a thorough characterization of each polymer by SEC coupled with an UV/Vis detector (Supplementary

Fig. 2b), which confirmed a successful conjugation and the absence of any unbound dye molecules. However, the introduction of charges by the dye significantly affected the self-assembly behaviour of the block copolymer (data not shown), in particular as the subsequent *ex vivo* experiments required high contents of labelled polymers (≥ 20 %). Consequently, only the spherical micelles could be obtained as a pure phase by this approach. Therefore, we also pursued a second strategy, where the dye labelling was performed subsequent to the assembly of the block copolymer P($BA_{50}$-*co*-$PDSA_5$)-*b*-$PEO_{125}$-$N_3$ (**P2**) and its crosslinking (experimental details are given in the chapter "nanostructure formulation").

*Experimental procedure for the synthesis of P($BA_{50}$-co-$PDSA_5$)-NHS (**P1**)*

In a microwave vial, BA (4.21 g, 32.8 mmol, 63.64 eq.), PDSA (0.79 g, 3.3 mmol, 6.36 eq.), PABTC-NHS (173.1 mg, 0.52 mmol, 1 eq.) were weighed and dissolved in 1,4-dioxane (0.64 mL). AIBN solution (218.3 µL of a 2 wt% solution in 1,4-dioxane, 25.8 µmol, 0.05 eq.) and 1,2,5-trioxane (~ 5 mg) were added, the vial was sealed with a rubber septum and the solution degassed under stirring by a stream of nitrogen for 20 min. Subsequently, a sample for $^1$H-NMR (100 µL, $CDCl_3$) was taken and the vial suspended in a preheated oil bath at 70 °C for 24 h. The conversion (79 %) was determined by $^1$H-NMR (50 µL, $CDCl_3$) by the ratio of the monomer vinyl signals to the 1,3,5-trioxane signal. After quenching by cooling, $Et_2O$ (2 mL) was added and the mixture precipitated in *n*-hexane (20 mL). The residual monomer was removed by repeated centrifugation (3 × 11000 rpm, 5 min), decantation and resuspension steps in *n*-hexane (10 mL). Afterwards the purified polymer was dried in a vacuum oven at 40 °C for 3 days. $^1$H-NMR (20 mg, $CDCl_3$) and SEC (10 mg, DMAc + 0.21% LiCl) was measured for characterization.

$^1$H-NMR ($CDCl_3$, 300 MHz): $\delta$ = 0.95 (3H, m, BA, -$CH_3$), 1.39 (2H, m, BA, -$CH_2$-), 1.61 (2H, m, BA, -$CH_2$-), 1.81-2.59 (3H, m, backbone, -$CH_2$-CH-), 2.84 (0.09H, s, NHS), 3.06 (0.2H, s, PDSA, -$CH_2$-S-S-), 4.05 (2H, s, BA, -O-$CH_2$-), 4.32 (0.2H, s, PDSA, -O-$CH_2$-), 7.17 (0.1H, s, PDSA, py), 7.77 (0.1H, s, PDSA, py), 8.49 (0.1H, s, PDSA, py).

SEC (eluent: DMAc + 0.21% LiCl, PMMA-standard): $M_n$: 8 100 g mol$^{-1}$, Đ = 1.13.

*Experimental procedure for the synthesis of P($BA_{50}$-co-$PDSA_5$)-b-$PEO_{125}$-$N_3$ (**P2**)*

At the start, P($BA_{50}$-*co*-$PDSA_5$)-NHS (**P1**) (1.2 eq., 1.27 g, 0.16 mmol) was dissolved in dry DMF (30 mL) under stirring. Separately, $N_3$-$PEO_{125}$-$NH_2$ (1 eq., 724 mg, 0.13 mmol) was dissolved in dry DMF (30 mL) by gentle heating and triethylamine (2 eq., 36.6 µL, 0.26 mmol) was added. The mixture of $N_3$-$PEO_{125}$-$NH_2$ and triethylamine was slowly added to the solution containing **P1** *via* a syringe pump (3 mL h$^{-1}$) under vigorous stirring at room temperature.

Subsequently, the reaction mixture was stirred at room temperature for 24 h. SEC (20 µL, DMAc + 0.21% LiCl, PMMA-calibration) was measured to check the successful coupling. After completion, the solvent was evaporated, the residue dissolved in acetone and precipitated in $Et_2O$:*n*-hexane 1:1 to remove the excess of P($BA_{50}$-*co*-$PDSA_5$)-NHS. After repeated centrifugation (3 x 11 000, 5 min) and decantation steps, the solvent was evaporated and the product dried in a vacuum oven at 40 °C overnight. SEC (10 mg, DMAc + 0.21% LiCl, PMMA-calibration) and $^1$H-NMR was measured after purification.

$^1$H-NMR ($CDCl_3$, 300 MHz): $\delta$ = 0.92 (3H, t, $^3J$, 6.0 Hz, BA, -$CH_3$), 1.24-1.77 (4H, m, BA, -$CH_2$-$CH_2$-), 1.82-2.46 (3H, m, backbone, -$CH_2$-CH-), 3.03 (0.2H, s, PDSA, -$CH_2$-S-S-), 3.64 (9.3H, s, mPEO), 4.02 (2H, s, BA, -O-$CH_2$-), 4.30 (0.2H, s, PDSA, -O-$CH_2$-), 7.13 (0.1H, s, PDSA, py), 7.73 (0.1H, s, PDSA, py), 8.48 (0.1H, s, PDSA, py).

SEC (eluent: DMAc + 0.21% LiCl, PMMA-standard): $M_n$: 18 300 g mol$^{-1}$, Đ = 1.14.

*Experimental procedure for synthesis of the polymer AF488-$PEO_{125}$-$NH_2$ (**P3**) via SPAAC*

$N_3$-$PEO_{125}$-$NH_2$ (20.7 mg, 3.78 µmol, 1 eq.) was dissolved in DMF (1 mL) and a solution of DBCO functionalized fluorescent dye AlexaFluor 488 (3 mg, 3.78 µmol, 1 eq.) in 1 mL DMF was added dropwise under stirring. The mixture was stirred at room temperature overnight to ensure a quantitative conjugation. Subsequently, SEC (20 µL, DMAc + 0.21% LiCl, UV/Vis detection 480-513 nm) was measured to analyze the conjugation reaction, the solvent evaporated and the product dried in a vacuum oven at 40 °C.

SEC (eluent: DMAc + 0.21% LiCl, PEG-standard): $M_n$: 5 400 g mol$^{-1}$, Đ = 1.07.

*Experimental procedure for the synthesis of P($BA_{47}$-co-$PDSA_5$)-NHS (**P4**)*

This second batch of the hydrophobic copolymer was prepared in a similar procedure as given for polymer **P1**. The following amounts were used: 4.21 g BA (32.8 mmol, 63.64 eq.), 0.79 g PDSA (3.3 mmol, 6.36 eq.), 173.1 mg PABTC-NHS (0.52 mmol, 1 eq.), 0.64 mL 1,4-dioxane, 218.3 µL of the AIBN solution (2 wt% solution in 1,4-dioxane, 25.8 µmol, 0.05 eq.). In this case an overall conversion of 73% was reached after 24 h.

SEC (eluent: DMAc + 0.21% LiCl, PMMA-standard): $M_n$: 8 100 g mol$^{-1}$, Đ = 1.13.

*Experimental procedure for the synthesis of P($BA_{47}$-co-$PDSA_5$)-b-$PEO_{125}$-AF488 (**P5**)*

The labelled block copolymer was prepared in a similar procedure as given for polymer **P2**. Instead of $N_3$-$PEO_{125}$-$NH_2$ the labelled polymer AF488-$PEO_{125}$-$NH_2$ (**P3**) was used for coupling. The following amounts were used: 32.2 mg P($BA_{47}$-*co*-$PDSA_5$)-NHS (**P4**) (1.2 eq., 4.54 µmol), 23.7 mg AF488-$PEO_{125}$-$NH_2$ (**P3**) (1 eq., 3.78 µmol), 0.77 g triethylamine (2 eq., 1.05 µL, 7.57 µmol), 10 ml DMF.

SEC (eluent: DMAc + 0.21% LiCl, PMMA-standard): $M_n$: 15 000 g $mol^{-1}$, Đ = 1.11.

***Nanostructure formulation***

The nanostructures were formulated *via* a solvent switch technique. First the polymer was dissolved in a non-selective co-solvent and water was slowly, but continuously added to reach a solvent ratio of 1:1 which triggered already the particle formation. The remaining co-solvent was removed by dialysis. Small spherical micelles with a hydrodynamic diameter ($D_H$) of ~ 25 nm were formed using acetone as co-solvent. Starting from a 1:1 mixture of DMSO and acetone wormlike micelles were obtained after switching the solvent to water, while pure DMSO resulted in the formation of pure vesicles ($D_H$ ~ 116 nm).

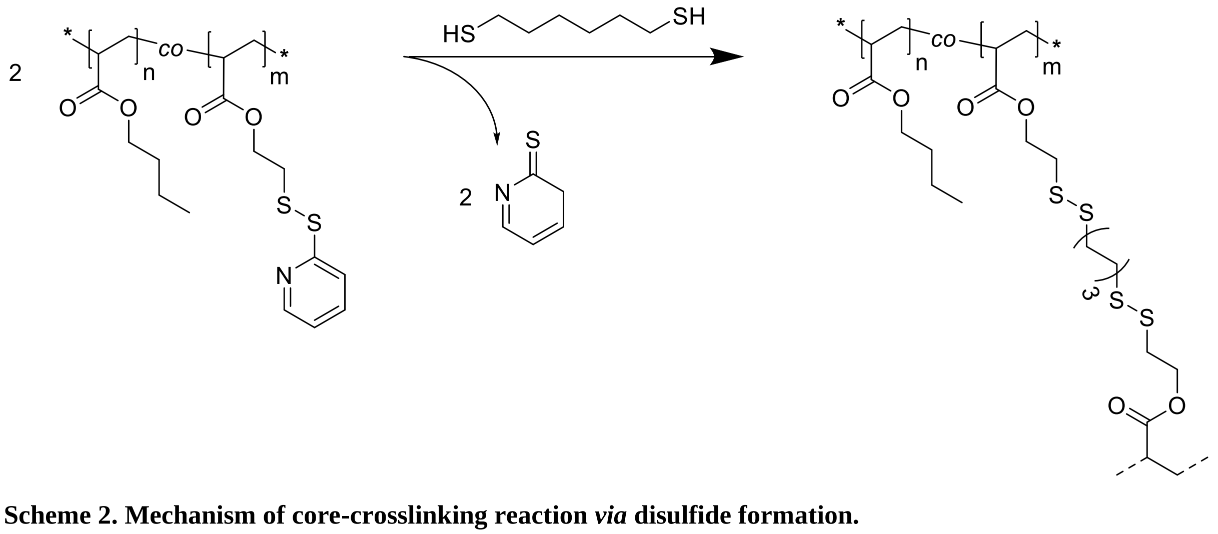


**Scheme 2. Mechanism of core-crosslinking reaction *via* disulfide formation.**

To stabilize the nanoparticles, the PDSA units were used to crosslink the obtained nanostructures by formation of disulfide links with added 1,6-hexanedithiol, which preferentially diffuses into the hydrophobic core of the micelles (Supplementary Scheme 2). The course of the reaction was monitored by UV/Vis absorption measurements (Supplementary Fig. 3), which prove the release of the by-product 2-mercaptopyridine (absorbance maximum at 343 nm). Control experiments without crosslinker and variations of the added equivalents of crosslinker verified that the crosslinking is efficient with addition of 0.5 equivalents of 1,6-hexanedithiol. The slight deviation from the maximal achievable release (in case of 1 or 2 eq. of added crosslinker) is related to a few unbound ends of the crosslinker, which occurs due to a limited mobility of the already mostly crosslinked chains within the core. Another prove for the efficient crosslinking is given by DLS measurements in the non-selective solvent THF before and after crosslinking (Supplementary Fig. 4). Moreover, no significant structural changes of the nanostructures were observed after crosslinking by DLS (Supplementary Fig. 5). Further information on the crosslinked nanostructures are summarized in Supplementary Table 2. The resulting nanostructures were characterized by DLS (Supplementary Table 3, Fig. 6).

**Table 2. Comparison of the wormlike micelles before and after core-crosslinking.**

| *Sample* | *Assembled from* | *$D_H$*[a] *[nm]* | *PDI*[a] |
|---|---|---|---|
| ***M3*** | Acetone/DMSO | 153.1 ± 3.7 | 0.26 ± 0.02 |
| ***M3-CL*** | Acetone/DMSO | 163.8 ± 2.9 | 0.25 ± 0.03 |

**a,** Determined by DLS measurements of the purified self-assembled structures (three technical replicates, c: 2 mg mL$^{-1}$).

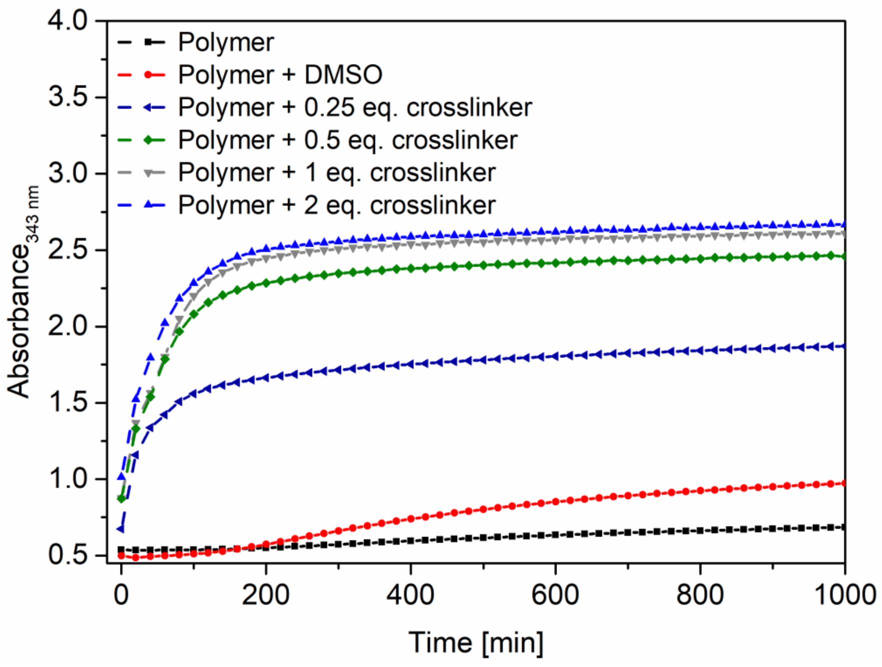

**Fig. 3 Kinetic study of crosslinking reaction *via* time dependent UV/Vis absorption measurements**. Absorbance of 2-mercaptopyridine by-product during disulfide formation recorded at 343 nm (Polymer: P($BA_{53}$-*co*-$PDSA_5$)-*b*-$PEO_{125}$, 2 mg $mL^{-1}$ in $H_2O$; crosslinker: HDT in DMSO, 1 mg $mL^{-1}$).

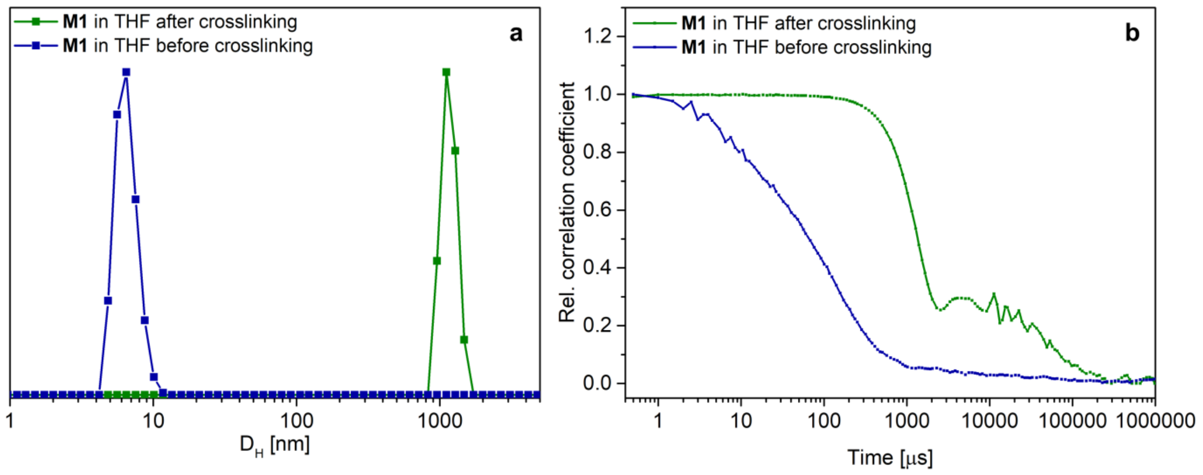


**Fig. 4 Characterization of the nanostructure core-crosslinking by DLS** (c: 1 mg $mL^{-1}$). **a,** Number-weight size distribution of sample **M1** in the non-selective solvent THF before and after core-crosslinking reaction. **b,** Correlation function of sample **M1** in the non-selective solvent THF before and after core-crosslinking reaction.

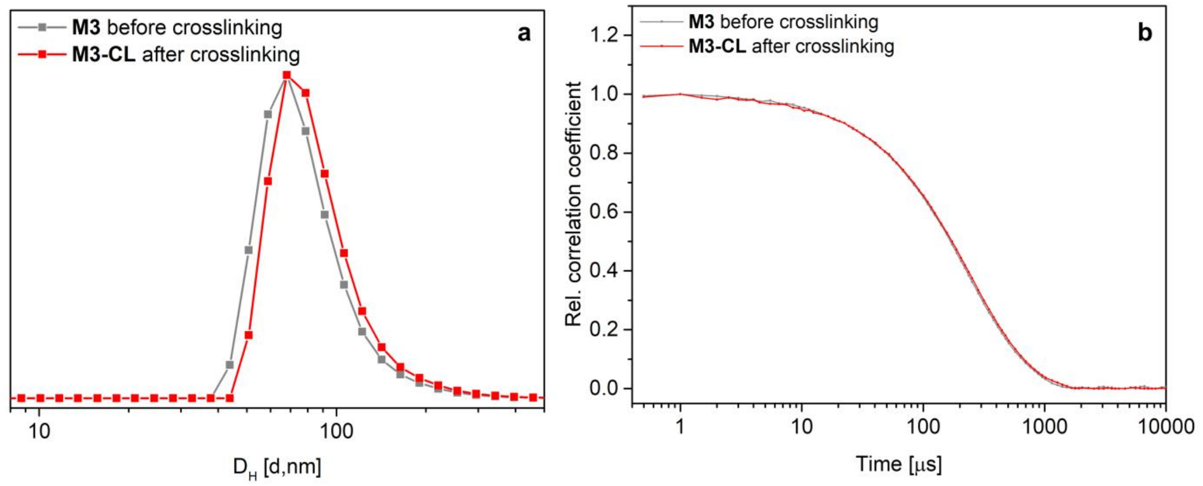


**Fig. 5 DLS characterization of wormlike micelles before and after core-crosslinking (**three technical replicates, c: 1 mg $mL^{-1}$). **a,** Number weight size distribution of **M3** before and after crosslinking. **b,** Correlation function of **M3** before and after crosslinking.

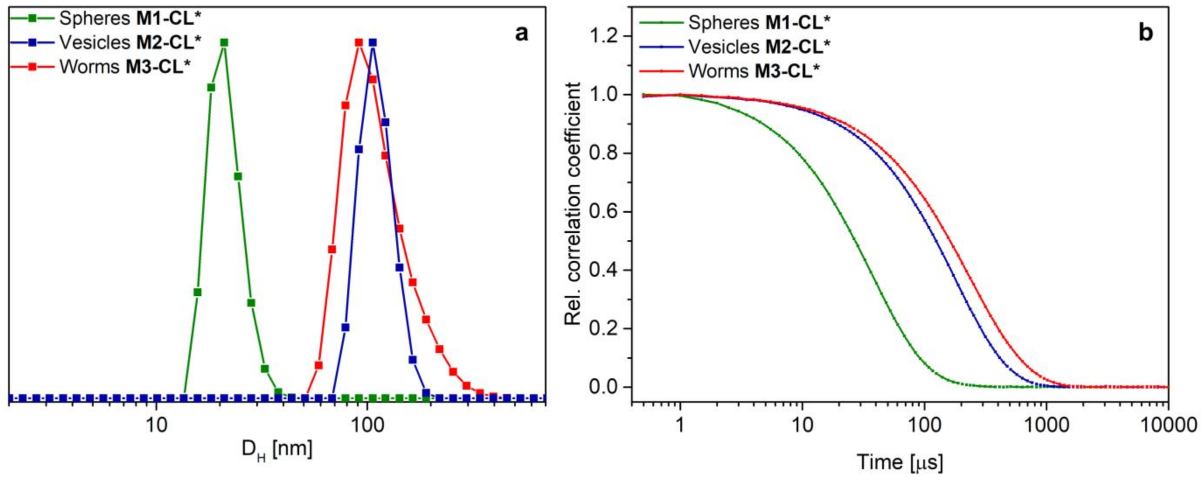


**Fig. 6 Characterization of the prepared nanostructures by DLS (**three technical replicates, c: 1 mg mL$^{-1}$). **a,** Number-weight size distribution of samples **M1-3-CL***. **b,** Correlation function of samples **M1-3-CL***.

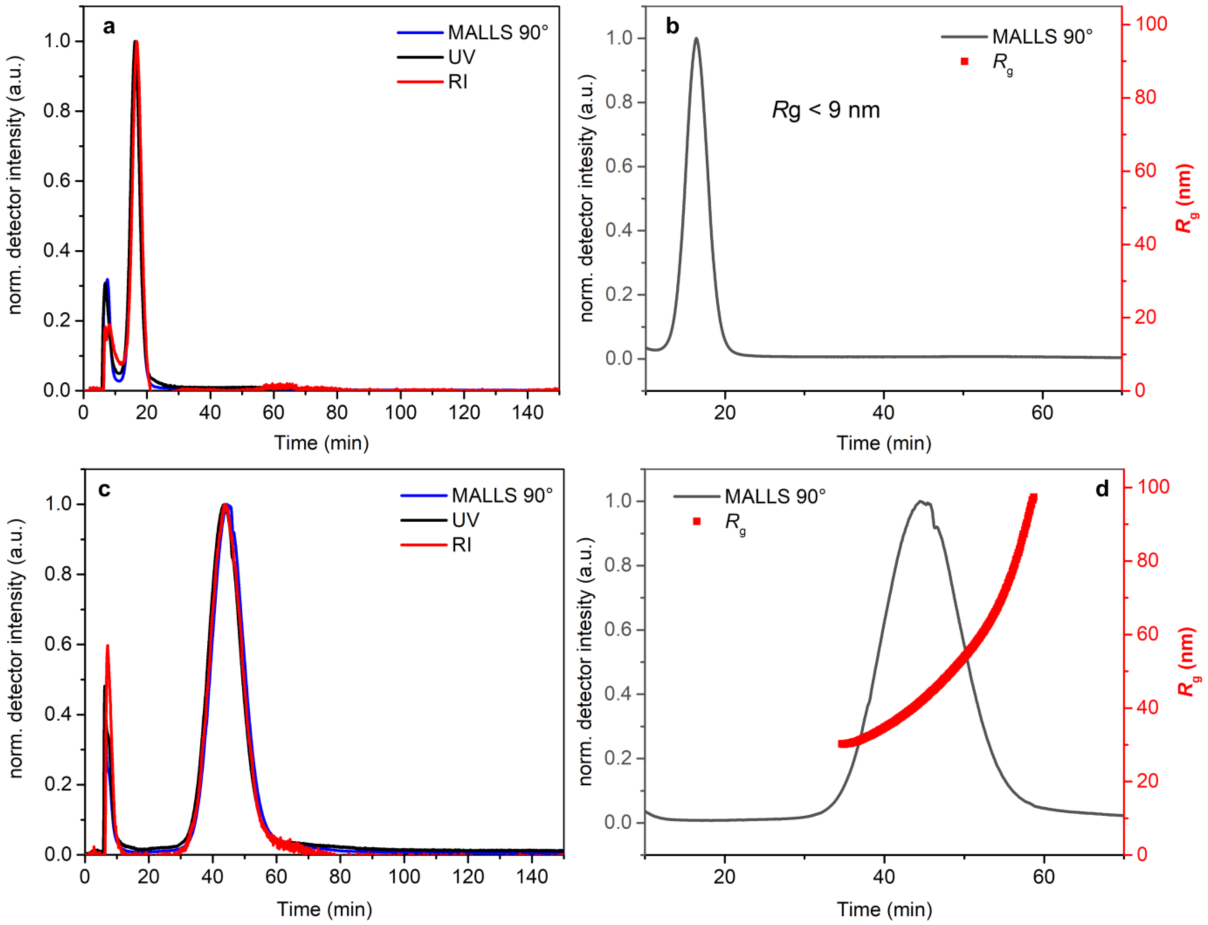

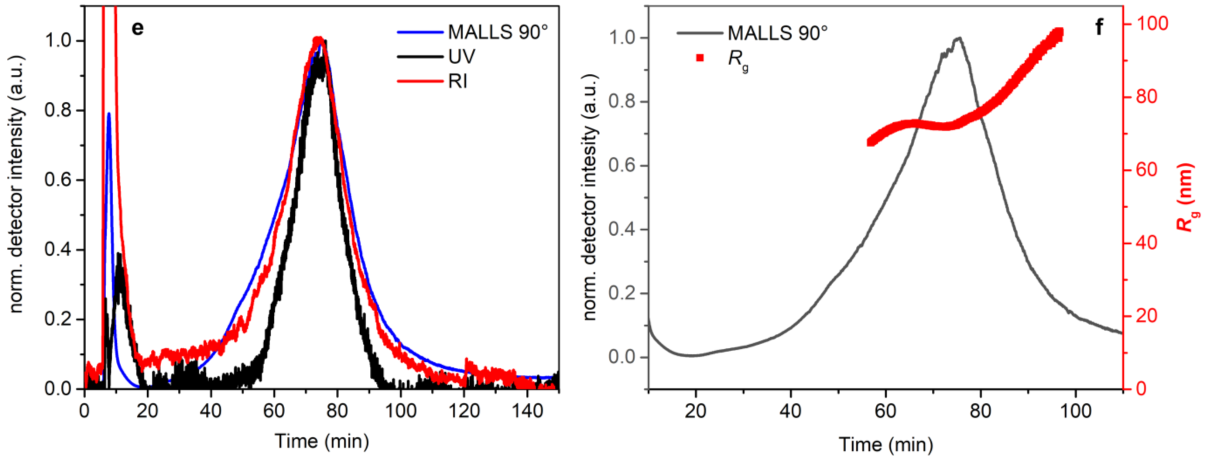


**Fig. 7 Normalized elugrams of core-crosslinked nanostructures by asymmetric flow field-flow fractionation (AF4). a,** Distribution of spheres **M1-CL*** ($c$ = 1.35 mg mL$^{-1}$) by RI, UV and MALLS detection. **b,** MALLS 90 °C scattering trace of **M1-CL*** and corresponding $R_g$ values. **c,** Distribution of vesicles **M2-CL*** ($c$ = 1.73 mg mL$^{-1}$) by RI, UV and MALLS detection. **d,** MALLS 90 °C scattering trace of **M2-CL*** and corresponding $R_g$ values. **e,** Distribution of worms **M3-CL*** ($c$ = 0.8 mg mL$^{-1}$) by RI, UV and MALLS detection. **f,** MALLS 90 °C scattering trace of **M3-CL*** and corresponding $R_g$ values.

**Table 3. Overview of the synthesized micelles including abbreviations and characterization.**

| *Sample* | *Morphology* | *Assembled from* | *$D_H$[a]* *[nm]* | *$D_C$[b]* *[nm]* | *PDI[a]* | *ζ-Potential[a]* *[mV]* |
|---|---|---|---|---|---|---|
| ***M1-CL**** | Spheres | Acetone | 25.6 ± 0.3 | 15.5 ± 2.3 | 0.10 ± 0.00 | - 10.5 ± 3.9 |
| ***M2-CL**** | Vesicles | DMSO | 116.1 ± 4.2 | 88.1 ± 16.4 | 0.04 ± 0.01 | - 23.6 ± 0.6 |
| ***M3-CL**** | Worms | Acetone/DMSO | 153.6 ± 1.2 | 20.5 ± 2.2 | 0.22 ± 0.01 | - 17.5 ± 0.3 |

**a,** Determined by DLS measurements of the purified self-assembled structures (three technical replicates, c: 1 mg mL$^{-1}$). **b,** Determined by graphical analysis of ≥ 100 particles in the cryo-TEM images.*Measured after core-crosslinking and labeling.

*General experimental procedure for the assembly of the nanostructures*

The block copolymer P(BA$_{50}$-*co*-PDSA$_5$)-*b*-PEO$_{125}$-N$_3$ **P2** (58.9 mg) was first dissolved in the respective organic solvent (acetone or DMSO) or the 1:1 solvent mixture (14.725 mL) and ultrapure water (14.725 mL) was added successively *via* syringe pump (1 mL h$^{-1}$) under vigorous stirring. Afterwards, the organic solvent was removed by dialysis against deionized water for 24 h (two solvent exchanges, MWCO: 3.5 - 5 kDa).

*General experimental procedure for the crosslinking of the nanostructures*

The structures were core-crosslinked by the addition of 1,6-hexanedithiol (1.658 mL of a 1 mg $mL^{-1}$ stock solution in DMSO, 0.5 eq. of PDSA groups) and stirring overnight at room temperature. Subsequently, the dispersion was purified from the organic solvent and any residual crosslinker by dialysis for 3 days (5 water exchanges MWCO: 3.5 - 5 kDa) against deionized water.

*Experimental procedure for the preparation of the labelled spherical micelles*

In case of the spherical micelles, the labelled nanostructures were directly prepared by assembling a mixture of the non-labelled block copolymer $P(BA_{50}$-*co*-$PDSA_5)$-*b*-$PEO_{125}$-$N_3$ (**P2**) (4 eq.) with the prelabeled polymer $P(BA_{50}$-*co*-$PDSA_5)$-*b*-$PEO_{125}$-AF488 (**P5**) (1 eq.) from pure acetone according to the procedure described above. The micelles were subsequently crosslinked by addition of 0.5 eq. of 1,6-hexanedithiol in accordance with the other samples. DLS indicated the formation of pure spherical micelles and the concentration was determined gravimetrically (n = 3) after lyophilization.

*Experimental procedure for the labeling of the pre-assembled and crosslinked nanostructures*

In case of the wormlike micelles and vesicles, the labeling reaction was performed by SPAAC after the crosslinking step. Therefore, DBCO functionalized Alexa Fluor 488 (699.61 µL of a 1 mg $mL^{-1}$ stock solution in $H_2O$, 0.2 eq. of azide groups) was added to the dispersion (2 mg $mL^{-1}$) and stirred at room temperature for at least 48 h. Any residual, uncoupled dye was removed by dialysis (MWCO: 3.5 - 5 kDa) first against sodium chloride solution (0.1 M) and then against water. The final concentration was determined gravimetrically (n = 3) after lyophilization.

### *Ex vivo experiments*

*Human biopsies*

The biopsies were collected from patients without and with inflammatory bowel disease (IBD) at the University Hospital of Jena. The study was approved by the committee of human ethics (3285-10/11). All participants were informed about the aim of the project and gave informed consent to participate in this study. The donors without IBD and any inflammation in gastrointestinal tract (GIT) were used as healthy controls. The state of inflammation in IBD patients was determined by endoscopic and histopathological features. Histological index score Nancy[3] (0-4) and Mayo score (0-3) were used for ulcerative colitis (UC), and simplified endoscopic activity score was calculated for the biopsied bowel segment in Crohn's disease (SES-CD, 0-12)[4] (Supplementary Table 4).

**Table 4. The degree of inflammation in IBD patients.**

| *Patient* | *1* | *2* | *3* | *4* | *5* | *6* | *7* | *8* | *9* | *10* | *11* | *12* | *13* | *14* |
|---|---|---|---|---|---|---|---|---|---|---|---|---|---|---|
| *IBD* | UC | CD | CD | UC | UC | UC | CD | UC | CD | UC | UC | CD | CD | UC |
| *NI/ Mayo-Score/ SES-CD* | 3/3 | 4/2 | 1/4 | 0/0 | 4/2 | 3/2 | 4/0 | 2/1 | 4/0 | 3/2 | 2/1 | 2/2 | 2/2 | 2/3 |

IBD - inflammatory bowel disease, UC - ulcerative colitis, CD - Crohn's disease, NI - histological index score Nancy (0-4), Mayo score (0-3), SES-CD - simplified endoscopic activity score for Crohn's disease (0-12).

The colon biopsies were obtained using 3.4 mm round Biopsy Forceps without spike (ENDO Flex) during routine endoscopy by experienced endoscopists. Biopsies were immediately transferred into previously oxygenated ice-cold modified Krebs-Ringer bicarbonate (mKRB) buffer and transported to the laboratory within 5-7 min. Biopsies were carefully unfolded under the Stereo Microscope Stemi 305 (Zeiss). Silicone rubber disks (190 µm thick) and thin plastic disks (220 µm) with a 2.5-mm centered hole were used in the adapters for the Ussing chamber (Warner Instruments, Inc., Hamden, CT, USA) to fix the tissue with Histoacryl Tissue Glue (BBraun) and prevent leakage of the nanoparticles through any damage of the biopsies' edges[5,6]. Buffer was immediately added to the luminal and mucosal part of the Ussing chambers.

*Ussing chamber technique to study translocation of nanoparticles*

The Ussing chambers were filled with modified Krebs-Ringer bicarbonate (mKRB) buffer (concentrations of ingredients in mM $L^{-1}$: $Na^+$ 140, $Cl^-$ 123.8, $K^+$ 5.4, $H_2PO_4^-$ 0.6, $HPO_4^{2-}$ 2.4, $Ca^{2+}$ 1.2, $Mg^{2+}$ 1.2, $HCO_3^-$ 21, D(+)-glucose 10.0, –OH-butyrate 0.5, glutamine 2.5, D(+)-mannose 10.0), which was continuously oxygenated (95% $O_2$ and 5% $CO_2$) and kept at a pH of 7.4 and a temperature of 37 °C by water circulating in reservoir water jacket[5,7]. The same buffer was used for calibrations and for transferring the biopsies to the laboratory (see above).

The mKRB buffer was replaced after 20-30 min of equilibration with 10 mL of fresh mKRB buffer on the mucosal part and 10 mL of mKRB buffer containing 100 µg $mL^{-1}$ of nanoparticles on the luminal part of the Ussing chamber using an integrated drainage system.

At the beginning of the experiment ($T_0$), after 1 h ($T_1$) and after 2 h ($T_2$) aliquots of buffer were collected from both the luminal (100 µl) and the mucosal part (500 µl) and immediately replaced with 400 µl of fresh buffer solution in the mucosal part. The concentrations of nanoparticles in these aliquots were estimated from the fluorescence intensity (recorded on a

Spectrofluorometer FP-8500, Jasco) of the solution in comparison with a previously recorded calibration curve (defined solutions of nanoparticles in the buffer). The determined average concentrations at each time point are given in Supplementary Table 5 for the respectively nanoparticles applied to healthy tissue or inflamed tissue from patients with IBD.

**Table 5. Time-dependent change of the concentration of micelles in luminal part.**

| | *healthy* | | | *IBD* | | |
|---|---|---|---|---|---|---|
| | $T_0$ (0 h) | $T_1$ (1 h) | $T_2$ (2 h) | $T_0$ (0 h) | $T_1$ (1 h) | $T_2$ (2 h) |
| *spherical micelles* | 100.0 | 89.83 ± 1.33*** | 82.43 ± 1.49*** § ß | 100.0 | 85.25 ± 4.49* | 78.37 ± 5.39** ‡ |
| *vesicles* | 100.0 | 96.29 ± 1.65 | 95.15 ± 1.85 § | 100.0 | 99.87 ± 1.63 | 94.25 ± 0.85** # ‡ |
| *filomicelles* | 100.0 | 95.42 ± 1.78 | 96.29 ± 1.71 ß † | 100.0 | 90.86 ± 3.58 | 84.15 ± 2.81** # † |

The results are presented as the percentage of the initial concentration (M ± SEM); IBD - inflammatory bowel disease. n = 6 (n – number of donors/patients). One-way analysis of variance (ANOVA) Dunnett's or Tukey's multiple comparison test (*), *p < 0.05, **p < 0.001, ***p <0.0001 and two-tailed unpaired Student's t-test (§,#,†,‡) were used (‡p < 0.05, †p < 0.01, §,#p < 0.001, ß p <0.0001). $T_{0\text{-}2}$ - incubation time (h).

Integrity and viability of the tissue were controlled by electrophysiological measurements. Voltage / Current Clamp for 6 Ussing Chambers (VCC, K. Mussler Scientific Instruments, Germany) was connected to the chambers *via* Ag/AgCl voltage and current electrodes and set of agar-salt bridges (4% agar in 3M KCl). Clean Ag wire electrodes were chlorided prior to their use in household bleach for 24–48 h for stable measuanemts[8]. The inherent fluid resistance and electrode potentials were evaluated every time before the biopsy was mounted to the chamber. Operation mode was changed from open circuit to voltage clamp for measuring the short circuit current ($I_{sc}$) during the experiments[9-11]. A 20 µA transepithelial bipolar current with pulse durations of 100 ms was applied to the tissue. No significant differences in the tissue resistances (R, $\Omega \cdot cm^2$) were observed for the tested healthy and inflamed tissue in the beginning of the experiment (Supplementary Fig. 8c).

Tissue viability was evaluated by adding 6.6 µM forskolin solution (Sigma Aldrich, UK) to the mucosal chamber after the experiment. Forskolin, the cAMP-dependent $Cl^-$ secretagogue, changes ion transport and as a result the electrophysiological parameters of the viable tissue[12,13]. Only biopsies with a potential difference (PD) ≤ -0.5 mV and a response to the forskolin (especially drop in PD) were included in analyses[8,12].

The translocation of the nanoparticles through the tissue was evaluated using an apparent permeability coefficient (Papp, cm/s) (Supplementary Fig. 8b). The apparent permeability coefficients were calculated using the equation (2):

$$Papp = (dQ / dt) / (C_0 \times A) \qquad (2)$$

where $dQ/dt$ is the transport rate, defined as the slope obtained from linear regression of nanoparticles transport; $C_0$ is the initial concentration of nanoparticles on the luminal side; $A$ is the surface area (0.049 $cm^2$).

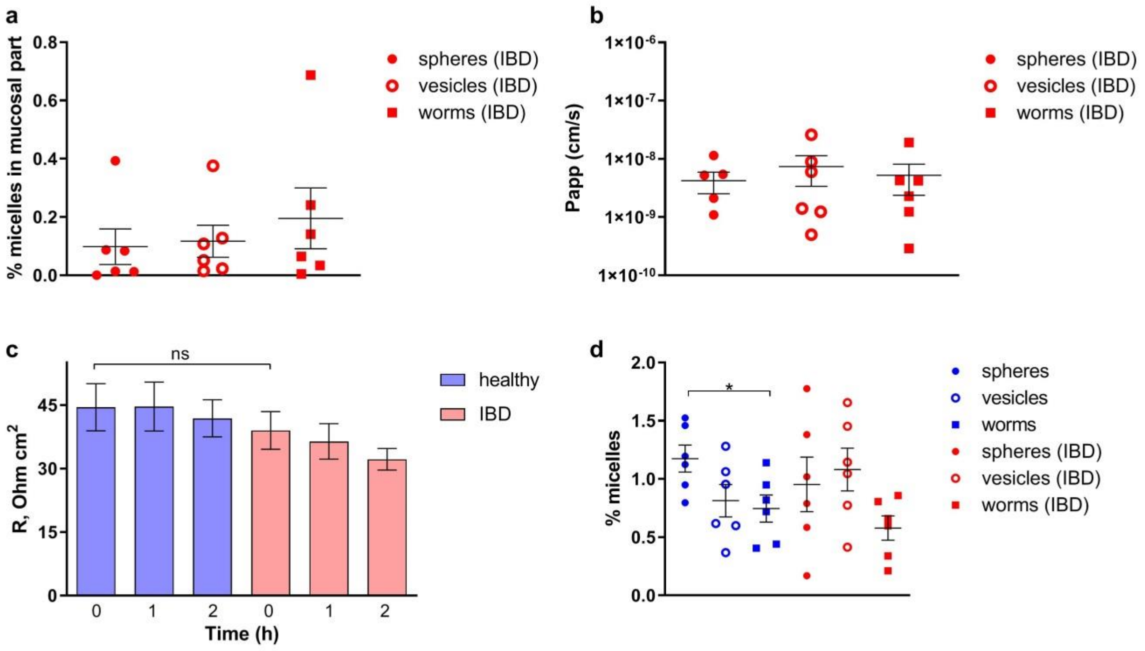


**Fig. 8 Additional information after Ussing chamber experiments. a,** ratio (in %) of translocated spherical micelles (full dots), polymeric vesicles (empty dots) and wormlike micelles (squares) through the inflamed mucosa. **b,** apparent permeability coefficient $P_{app}$ cm/s for the corresponding nanoparticles calculated for an exposed tissue area of 4.9 $mm^2$. **c,** The tissue resistance (R, $\Omega \cdot cm^2$) in the Ussing chamber experiments for healthy (blue) and inflamed (red) biopsies. **d,** The amount of nanoparticles washed off the human mucosa after Ussing chamber experiments. The results are presented as M ± SEM; n = 6 (n – number of donors/patients). Two-tailed unpaired Student's t-test (*) was used; *$p < 0.05$, ns - not significant.

We further verified that none of the three types of nanoparticles stick to the surface of Ussing chambers or are technically hindered to diffuse into the mucosal chamber by applying the nanoparticle solutions to empty Ussing chambers without tissue (Supplementary Fig. 9). No particle loss is observed after 2 h and equal amounts were found in the respective chambers as expected for an unhindered diffusion.

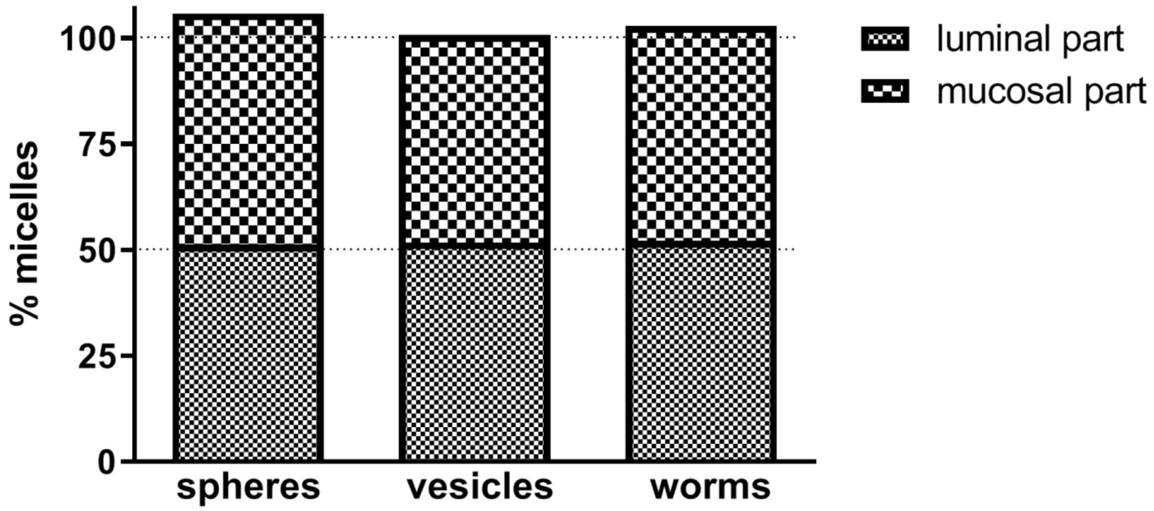


**Fig. 9 The distribution and content of nanoparticles in Ussing chambers without tissue after 2 h**. Nanoparticles were injected in luminal part of the chamber without tissue, while buffer was added at the same time to the mucosal part

*Localisation of the nanoparticles*

The biopsies were removed from the Ussing chambers after experiments and carefully washed with mKRB for determination of any nanoparticles attached to the surface of the tissue (Supplementary Fig. 8d). Subsequently the biopsies were fixed by placing them for 30 min in 4% paraformaldehyde with 25 mM glycine at room temperature and replace the solution with 15% and 30% sucrose solution (4°C) for overnight. Biopsies were then frozen in Tissue-Tek OCT Compound (Sakura) and sectioned into slices of 6 µm thickness by a cryotome (Leica CM 1050). An antibody for E-cadherin labelled with Alexa Fluor 647 (1:100, BD Biosciences), an antibody for CD11b labelled with Alexa Fluor 594 (1:300, BioLegend), and DAPI Fluoromount-G (SouthernBiotech) were used to selectively stain the tight junctions, the immune cells, and the nuclei of the cells, respectively. Biopsies that were not treated with nanoparticles, but still kept for the same time in Ussing chambers were used as controls. The localisation of the particles in correlation with the above-mentioned selective stains was observed with an Axio Observer Z.1 Microscope (Zeiss). Z-stacks (Supplementary Videos 1-6) and images were obtained using a Plan-Apochromat 40x/0.95 Korr M27 objective. Data analyses were performed using the Zeiss ZEN 2.3 (blue edition) software.

Microscopic images of the localisation of nanoparticles in healthy and inflamed human mucosa given in Fig.3 of the main manuscript are presented in Supplementary Fig. 10 and Fig. 11. Extended images of the colocalisations of particles and immune cells given in Fig. 4 of the main manuscript are presented in Supplementary Fig. 12.

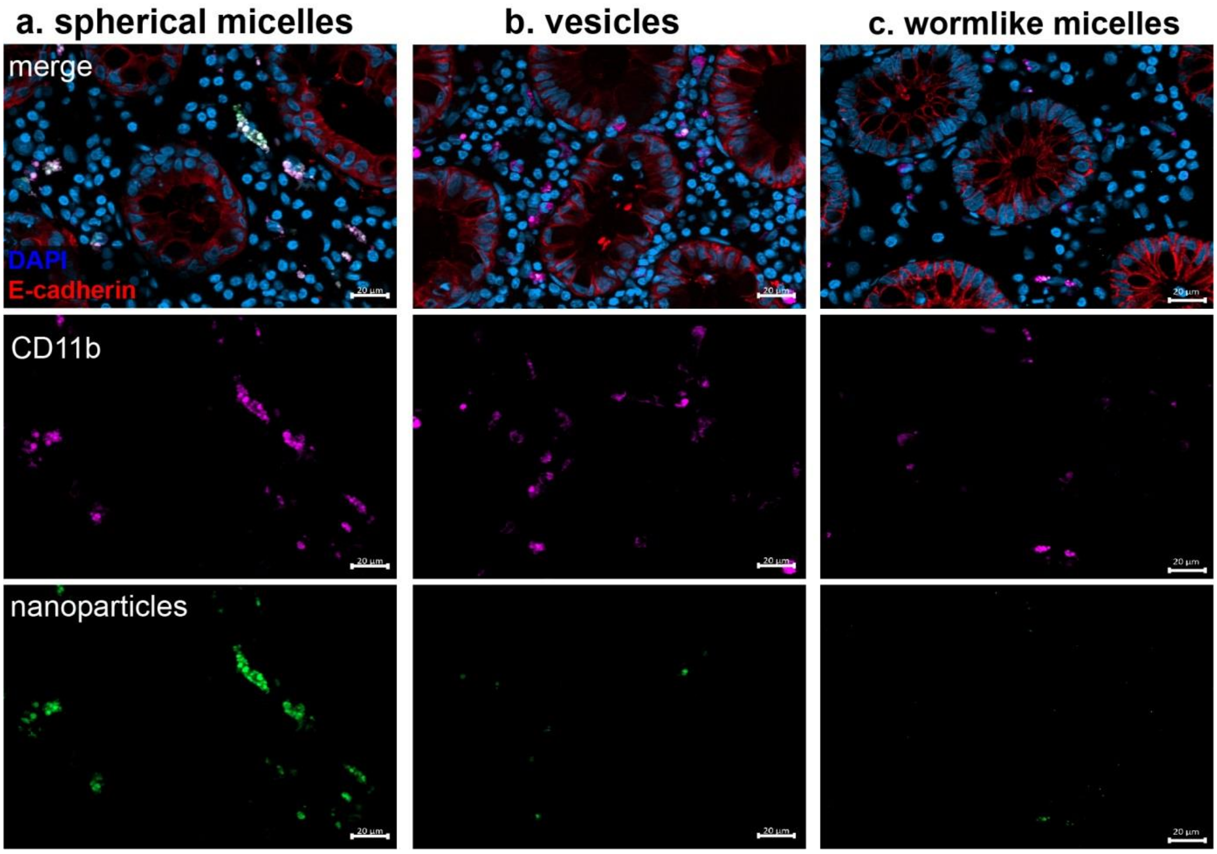


**Fig. 10 Localisation of nanoparticles in healthy human mucosa. a,b,c,** Localisation of spherical micelles (**a**), polymeric vesicles (**b**) and wormlike micelles (**c**) in healthy noninflamed human mucosa after 2 h incubation in Ussing chambers (green: nanoparticles; blue (DAPI): nuclei; red: E-cadherin (epithelium); violet: CD11b (immune cells)). Biopsies were sectioned (6 µm) by cryotome slicing. Scale bar is 20 µm.

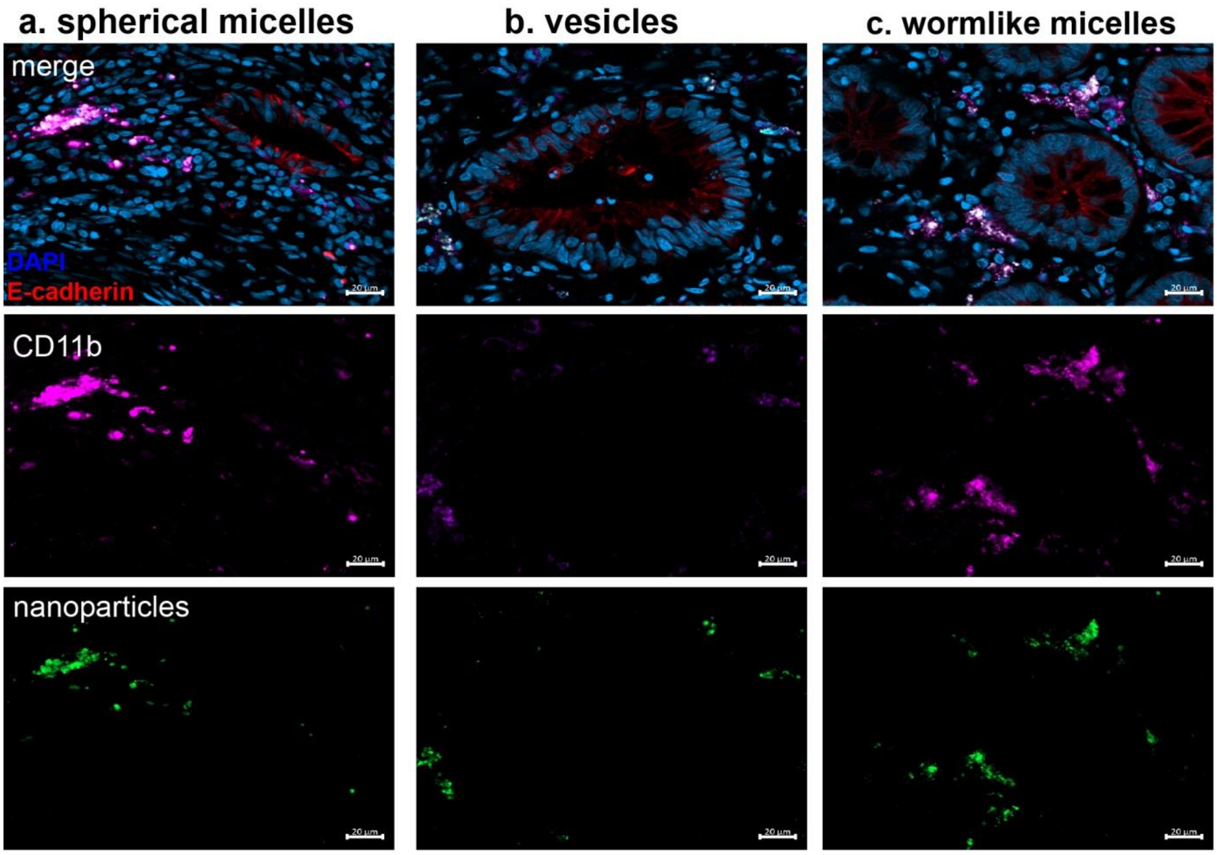


**Fig. 11 Localisation of nanoparticles in inflamed human mucosa. a,b,c,** Localisation of spherical micelles (**a**), polymeric vesicles (**b**) and wormlike micelles (**c**) in inflamed human mucosa after 2 h incubation in Ussing chambers (green: nanoparticles; blue (DAPI): nuclei; red: E-cadherin (epithelium); violet: CD11b (immune cells)). Biopsies were sectioned (6 µm) by cryotome slicing. Scale bar is 20 µm.

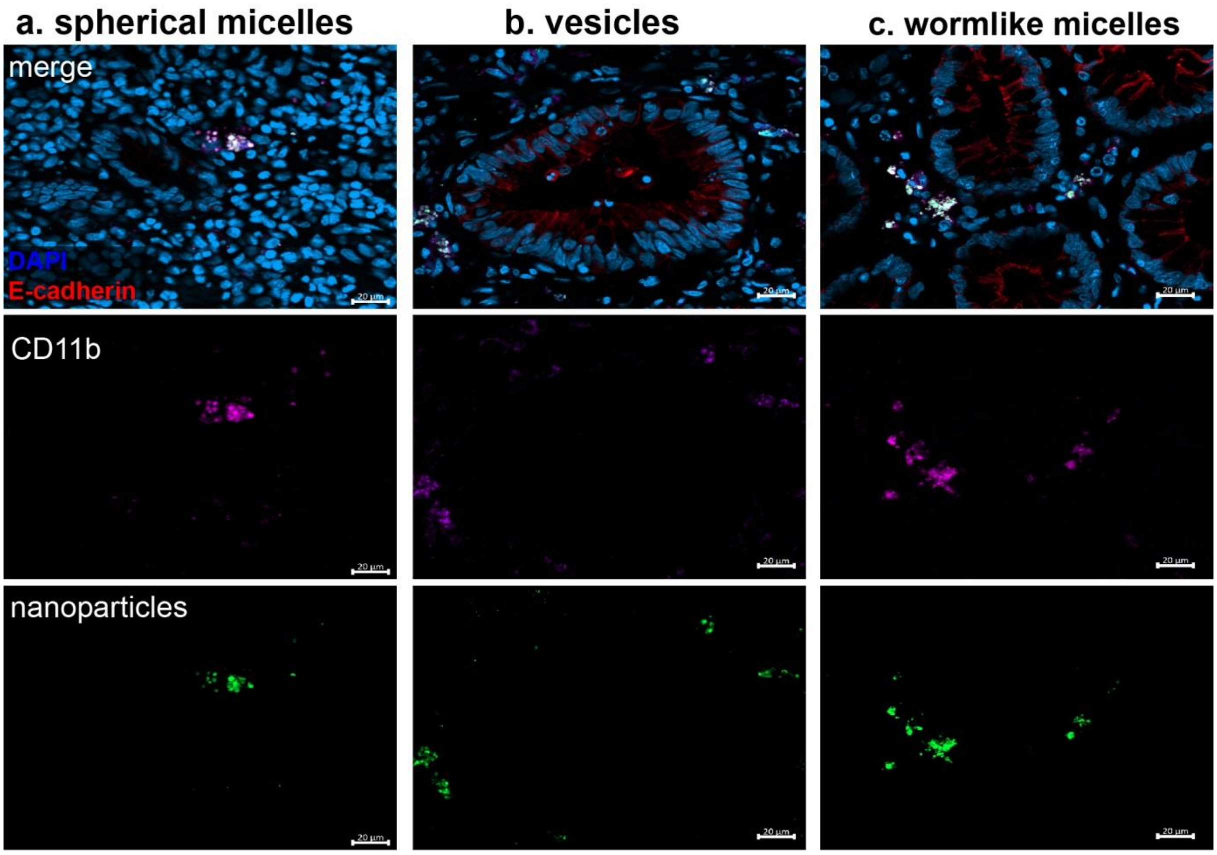


**Fig. 12 Accumulation of micelles in immune cells of inflamed human mucosa. a,b,c,** Accumulation of spherical micelles (**a**), vesicles (**b**) and wormlike micelles (**c**) in immune cells CD11b (violet:), (green: nanoparticles; blue (DAPI): nuclei; red: E-cadherin (epithelium)). Biopsies were sectioned (6 µm) by cryotome slicing. Scale bar is 20 µm.

*Statistical analysis*

Statistical analysis was performed using GraphPad Prism 9.0.0 (GraphPad Software, San Diego, California USA). For multiple comparisons the one-way analysis of variance (ANOVA) followed by Dunnett's or Tukey's tests were used. To compare two groups the two-tailed unpaired Student’s t-tests were performed. The results are presented as the mean value ± standard error of the mean (M ± SEM). Differences with $p = 0.05$ were considered statistically significant. The hydrodynamic diameters ($D_H$), PDI, ζ-Potentials are presented as the mean value ± standard deviation of the mean (M ± SD) of three technical replicates.
